# Genomic mating in outbred species: predicting cross usefulness with additive and total genetic covariance matrices

**DOI:** 10.1101/2021.01.05.425443

**Authors:** Marnin D. Wolfe, Ariel W. Chan, Peter Kulakow, Ismail Rabbi, Jean--Luc Jannink

## Abstract

Diverse crops are both outbred and clonally propagated. Breeders typically use truncation selection of parents and invest significant time, land and money evaluating the progeny of crosses to find exceptional genotypes. We developed and tested genomic *mate* selection criteria suitable for organisms of arbitrary homozygosity level where the full-sibling progeny are of direct interest as future parents and/or cultivars. We extended cross variance and covariance variance prediction to include dominance effects and predicted the multivariate selection index genetic variance of crosses based on haplotypes of proposed parents, marker effects and recombination frequencies. We combined the predicted mean and variance into usefulness criteria for parent and variety development. We present an empirical study of cassava (*Manihot esculenta*), a staple tropical root crop. We assessed the potential to predict the multivariate genetic distribution (means, variances and trait covariances) of 462 cassava families in terms of additive and total value using cross-validation. We were able to predict all genetic variances and most covariances with non-zero accuracy. We also tested a directional dominance model and found significant inbreeding depression for most traits and a boost in total merit accuracy for root yield. We predicted 47,083 possible crosses of 306 parents and contrasted them to those previously tested to show how mate selection can reveal new potential within the germplasm. We enable breeders to consider the potential of crosses to produce future parents (progeny with excellent breeding values) and varieties (progeny with top performance).

**Author Summary:** Breeders typically use truncation selection and invest significant resources evaluating progeny to find exceptional genotypes. We extended genetic variance and trait covariance prediction to include dominance and predicting the multivariate selection index variance. We enable mate selection based on potential to produce future parents (progeny with excellent breeding values) and/or varieties (progeny with top performance). Using cross-validation, we demonstrate that genetic variances and covariances can be predicted with non-zero accuracy in cassava, a staple tropical root crop.

## Introduction

Diverse crops ranging from staples (e.g., cassava, potato) to cash crops (e.g., cacao) to forestry products (e.g. eucalyptus) are both outbred and clonally propagated (Gemenet and Khan 2017). In these crops, exceptional genotypes can be immortalized and commercialized as clonal varieties. Few clonal crops are also inbred thus, like livestock, each cross segregates phenotypically to different degrees. Unlike seed crops (e.g. maize, wheat), inbreeding is unnecessary for product development.

Consider a breeding program implementing some form of genomic selection (GS) (Heffner *et al.* 2009; Jannink *et al.* 2010) on a population. All extant members and future progeny are or will be genotyped using genome-wide markers. Field evaluations are conducted at species- and trait-appropriate stages for one or more traits, on at least a subset of the genotypes. Genomic prediction is used to increase selection intensity and decrease cycle times by providing selection criteria for more genotypes, faster than would have otherwise have been possible (Hickey *et al.* 2017). The breeding scheme can be further divided in two parts (Gaynor *et al.* 2017; Santantonio and Robbins 2020; Werner *et al.* 2020) consisting of (1) population improvement by recurrent selection **(RS)** and (2) a variety development pipeline **(VDP).** RS is done in order to manage and improve the frequency of beneficial alleles in the population over time. The VDP consists of a series of field trials in which candidates’ performance is evaluated. For clonal crops, germplasm are advanced from one VDP stage to the next by vegetative propagation.

### The importance of matings and the need for mate selection criteria

Every cross is important. Crosses imply an opportunity and a risk. New matings generate genetic variation, the substrate on which selection can operate. However, for a breeder, new crosses require investment of time, land and money, especially considering the added costs of genotyping. Moreover, crosses may exhibit inbreeding depression or heterosis. Thus, matings serve the multiple purposes of producing new candidate breeding parents for RS and/or cultivars for VDP, potentially evaluated for multiple product profiles characterized by unique selection indices (SI).

Selection to drive improvement in the population’s mean over time to meet the objective of RS centers on allele-substitution effects and the breeding value (BV). For the VDP, selecting clones to advance for testing should be based on the total genetic value (TGV) of an individual which includes non-additive genetic effects such as dominance.

### Genomic predictions incorporating non-additive effects

Non-additive effects can be included in genomic predictions in a number of ways (Vitezica *et al.* 2013; Varona *et al.* 2018). Most of the literature so far has dealt with including non-additive effects in the prediction of the genetic values of an existing pool of selection candidates (Varona et al. 2018). Non-additive predictions have often been shown to increase prediction accuracy (Heslot *et al.* 2012; Wolfe *et al.* 2016b; Werner *et al.* 2020). In addition, the mean performance (mean TGV) of the progeny can deviate from the prediction based on the mean BV of the parents in the presence of non-additive effects. Genomic predictions of cross mean TGV have been applied to hybrid performance (GCA-SCA: (Alves *et al.* 2019) and mate allocation (Toro and Varona 2010). Predictions can also include genome-wide inbreeding/overdominance effects, also referred to as directional dominance; (Xiang *et al.* 2016) and this has recently been shown to be advantageous in a simulated two-part clonal crop breeding scheme (Werner *et al.* 2020).

### Genomic mate selection for outbred, clonal crops

When one or both parents are heterozygous, offspring are expected to segregate for their BV and TGVs. The relative advantage of possible pairwise matings can best be distinguished when predictions of both the genetic mean *and* variance are available. The usefulness criterion (UC) or simply “usefulness” of a cross is a prediction of the mean performance of the selected superior fraction of the progeny: *UC* = *μ* + *i* × *σ* where *σ* is the predicted genetic standard deviation of the progeny and *i* is the standardized selection intensity (Zhong and Jannink 2007; Segelke *et al.* 2014; Lehermeier *et al.* 2017b). The additive genetic variance of an infinite pool of progeny from a cross can be predicted deterministically using the combination of genome-wide marker effects, a genetic map, and phased parental haplotypes (Lehermeier *et al.* 2017b). Until now, this approach has exclusively been applied to the prediction of additive genetic variance and covariance; (Neyhart *et al.* 2019). While (*Bijma et al.* 2020) predicted the gametic variance (rather than progeny variance) for outbred populations, most other applications are predictions of the variance of inbred lines derived from inbed founders (Zhong and Jannink 2007; Lehermeier *et al.* 2017b; Neyhart *et al.* 2019; Neyhart and Smith 2019; Allier *et al.* 2019a).

### Criteria and Methods Developed in this Article

In this article, we extend the deterministic prediction of progeny variances in several ways to maximize the utility and practicality of implementing genomic mate selection. First, we show how to include dominance in the prediction of cross genetic variance and we do so for founders of arbitrary inbreeding level. Next, we distinguish two types of cross usefulness: usefulness for recurrent selection (i.e. the predicted mean BV of offspring selected as parents; *UC_parent_*) and usefulness for variety development (i.e. the predicted mean TGV of clones advanced as varieties in the VDP; *UC_variety_*). Finally, since matings are usually chosen on the basis of multiple traits, we extend the prediction of cross variance to selection indices (SI). We show that to predict index variance, we must predict the full matrix of trait genetic variances and covariances, which can be done following (Neyhart *et al.* 2019; Allier *et al.* 2019a). We implement the core functions for multi-trait prediction of outbred cross variances including additive and dominance effects in an R package predCrossVar (https://wolfemd.github.io/predCrossVar/).

### Empirical Study of Cassava

We present an empirical study of the accuracy for predicting additive and non-additive genomic mate selection criteria. We set up a cross-validation scheme that measures the accuracy of predicting means, variances and usefulnesses of previously untested crosses using data from a real cassava *(Manihot esculenta)* breeding program. Cassava is one of the most important tropical staple foods, especially in Africa (http://faostat.fao.org). Among outbred, clonal crops, GS is relatively mature in cassava breeding (Yonis *et al*.; de Oliveira *et al.* 2012; Ly *et al.* 2013; Wolfe *et al.* 2016a; b, 2017; Okeke *et al.* 2017; Elias *et al.* 2018; Ozimati *et al.* 2018) because of the Next Generation Cassava Breeding Project (http://www.nextgencassava.org, est. 2012), and the species can serve as a model for many others. We leverage a validated GS pedigree (3199 clones, representing 4 generations, 209 parents, 462 full-sib families) with genome-wide phased haplotypes and a genetic map detailed in (Chan *et al.* 2019). We also compare the standard additive-dominance model in which dominance effects have a mean of zero to a directional dominance model (Xiang *et al.* 2016), which allows us to make first-time estimates of genome-wide inbreeding (homozygosity) effects. Finally, we report our empirical study in a fully reproducible (https://github.com/wolfemd/PredictOutbredCrossVar/) and documented (https://wolfemd.github.io/PredictOutbredCrossVar/) framework.

## Methods

### Formulation of Genomic Predictions and Selection Criteria

Below, we describe predictions that are applicable as selection criteria, first for genomic truncation selection *G_TS_* followed by extensions that enable mate selection *G_MS_*. Throughout, we distinguish selection criteria based on their suitability for evaluating the potential of individuals (for *G_TS_*) or crosses (for *G_MS_*) for RS vs. VDP.

#### *G_TS_*: Selecting genotypes with predictions about generation *t*

Genomic recurrent TS (*G_TS_*) evaluates existing individuals, either for their potential as parents (without regards to specific mates) and/or their potential as clonal cultivars.

Under a non-epistatic model, the total genetic values of individuals in the current population (time t) can be partitioned into a breeding value (*g^BV^*) and a dominance deviation *g^DD^*.

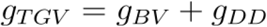

Consider a diploid population with n individuals genotyped at *p* biallelic genomic loci.

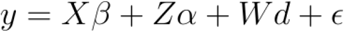

In this linear model, the *n* × 1 vector of phenotypic observations, *y* is modelled according to a combination of genetic and non-genetic effects. Fixed experimental design-related effects estimates are given by *β* ([*n* × *N_fixed_*]) and its corresponding incidence matrix *X*, where *N_fixed_* is the number of fixed factors. The elements of the [*n* × *p*] matrices *Z* and *W* contain column-centered marker genotypes:

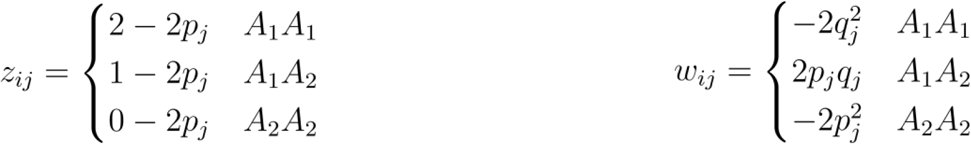

This encoding of genotypes results in marker effects (*α* and *d*) that correspond to allele substitution and dominance deviation effects (Vitezica *et al.* 2013). The marker effects can then be used to predict genomic estimated total genetic values 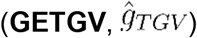 as the sum of the genomic estimated breeding value 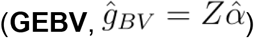 and a corresponding dominance deviation 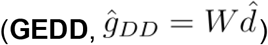.

The GEBV predicts the mean of offspring of a clone mated at random and as such is suitable for truncation RS of parents. The GETGV predicts the performance of each clone, rather than any property of its offspring and is useful for selection for variety advancement.

#### *G_MS_*: Selecting crosses with predictions about generation *t*+1

GEBV and GETGV enable us to do truncation selection. In order to implement mate selection, criteria that distinguish crosses are needed.

Progeny of crosses may segregate for both their breeding and total genetic values. Crosses may thus differ in their likelihood of producing progeny that are superior varieties (high (*g_TGV,t_*+1) and/or parents (high *g_BV,t_*+1). We focus here on distinguishing the best crosses on the basis of both their predicted genetic means *and* variances.

| Parameter | Breeding Values (BV) | Total Genetic Values (TGV) |
| --- | --- | --- |
| Mean | $\mu_{BV}$ | $\mu_{TGV}$ |
| Variance | $\sigma_{BV}^2$ | $\sigma_{TGV}^2 = \sigma_{BV}^2 + \sigma_{DD}^2$ |

##### Predicted cross means

The family mean can be predicted as the mean of parental breeding values.

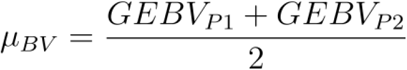

Dominance deviation can be included in order to predict the mean TGV (Toro and Varona 2010; Varona *et al.* 2018); see also (Falconer and Mackay 1996; Werner *et al.* 2020).

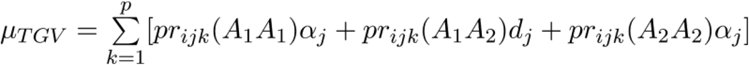

Here *pr_ijk_*(*A*_1_ *A*_1_), *pr_ijk_*(*A*_1_ *A*_2_) and *pr_ijk_*(*A*_2_ *A*_2_) represents the expected frequencies of each possible genotype in offspring of the *i*th and *j*th individual, and the summation, index by *k* is across *p* markers.

##### Predicted cross variances

(Lehermeier *et al.* 2017b) showed that the within-cross additive genetic variance can be predicted deterministically, as applied to the special case where progenitors are inbred and the terminal generation of progeny being predicted is also inbred (e.g. doubled-haploids or *S*_6_; see also (*Segelke *et al.* 2014; Neyhart *et al.* 2019; Allier et al.* 2019a). In addition, (Bijma *et al.* 2020) described the application of Lehermeier et al.’s approach to the prediction of additive *gametic* variance and selection of parents in animals. (*Lehermeier et al. 2017b; Bijma et al. 2020)* rely on the formula for the genetic variance under linkage disequilibrium, as given by (Lynch and Walsh 1998), Eqn. 5.16a.

Below, we show that a simple extension to predict dominance variance can be made using (Lynch and Walsh 1998), eqn. 5.16b. We predict both additive and dominance variances in an infinite population of full-siblings derived from outbred progenitors (i.e. parents with any inbreeding level).

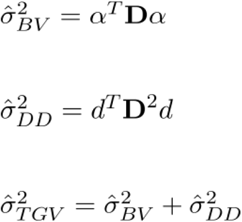

The *p* × *p* variance-covariance matrix, **D**, is the expected linkage disequilibrium among full-siblings by considering the expected pairwise recombination frequency and each parent’s haplotype phase.

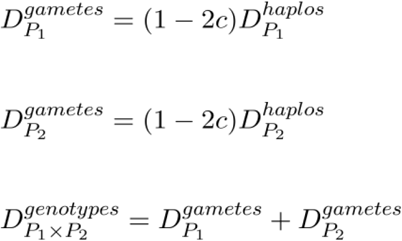

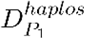 and 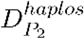 are simply the *p* × *p* covariance matrices associated with each parent’s respective 2 × *p* haplotype matrix, where elements are 1 if the counted allele is present, 0 otherwise. The *p* × *p* pairwise recombination frequencies matrix is c and can be derived from a genetic map. 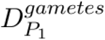 and 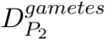 are the covariance matrices for each parents pool of possible gametes, whose covariances sum to give the expected covariances genotypes in the cross, 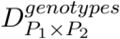. The genetic variances 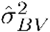 and 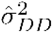 are thus predicted as above by using 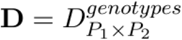.

##### Usefulness Criteria (UC) - Mean of superior family members

Given that predictions of genetic means and variances for a cross are available, they can be combined into a single cross selection criterion. We focus here on the **usefulness criterion (UC)**, which predicts the mean (breeding value) of the superior progeny from a cross, i.e. the mean *after* selection (Schnell and Utz 1976; Zhong and Jannink 2007; Lehermeier *et al.* 2017b). We note that predictions of cross means and variances may be used in other ways (e.g. (Bijma *et al.* 2020), but focus on UC. The *UC* = *μ* + *i* × *σ*, where *μ* is the predicted mean of the cross, *i* is the standardized within-family selection intensity and *σ* is the predicted cross standard deviation.

In the context of the two-part breeding scheme for GS in clonal crops, crosses may be *useful* either for producing new parents and/or new varieties. Therefore, we define two UCs:

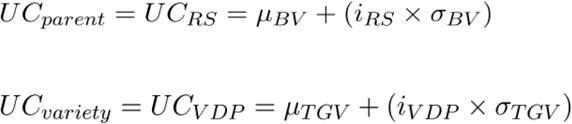

Notice that in addition to separate predictions of mean and variance for *UC_parent_* versus *UC_variety_*, two-part GS implies that the within-family intensity of selection for RS does not necessarily equal that of the VDP (see Santantonio and Robbins 2020).

#### Extension to multi-trait selection indices

Real selection is usually done based on the net merit of candidates relative to a multi-character selection index (SI). It is thus essential to consider multiple traits. Moreover, cross usefulness might be considered relative to multiple product profiles with differing likelihood of producing progeny with good multivariate genetic values, composed as a linear SI. Indeed, manipulating multivariate genetic variation is the primary advantage of recombination. Previous work on the UC and the prediction of cross-variance has considered single traits at a time (Schnell and Utz 1976; Zhong and Jannink 2007; Lehermeier *et al.* 2017b). (Neyhart *et al.* 2019) showed how to predict the correlated response to selection on one trait, using the predicted additive genetic covariance between traits (see also (Allier *et al.* 2019a). Using both the predicted variances and covariances among SI-component traits, we can therefore predict the mean and variance of a cross on the SI as follows:

1. Predict (co)variances for all traits on SI: Consider an index with two traits, *T*1 and *T*2.

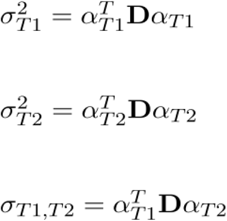 Apply to dominance by substituting n with *d* and squaring D.
2. Compute the predicted mean and variance on the selection index:

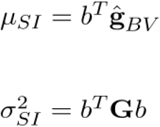

The *n* × *T* matrix 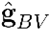 contains the GEBV for each trait and the *Tx*1 vector *b* are the index weights. The *T* × *T* matrix **G** is the additive (or total) genetic variance-covariance matrix for traits on the index.

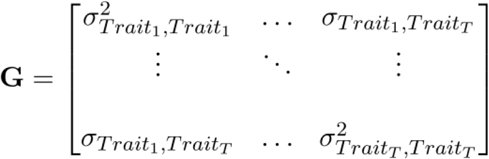

Based on these predictions of family means, variances and trait-covariances, we can finally compute the mean of selected family members on the index (i.e. the *UC_SI_*).

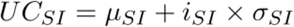

#### Including directional dominance as a genome-wide inbreeding effect

Many outbred, clonal crops are known to suffer from inbreeding depression. The typical genome-wide regression models the marker effects as drawn from a normal distribution, with mean zero and an estimated variance parameter. To include directional dominance, we model the genome-wide proportion of loci that are homozygous, f:

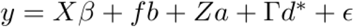

Xiang *et al.* (2016) showed that the effect estimated by *b* is the linear effect of overall homozygosity, interpreted as inbreeding depression or heterosis depending on its direction relative to each trait. The effect of over/under-dominance measured by *b* can be incorporated into the predicted means and variances by dividing *b* by the number of effects (*p*) and subtracting that value from the vector of dominance effects, to get 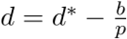 (Xiang *et al.* 2016; Varona *et al.* 2018; Werner *et al.* 2020). It is important to note that the partition of genetic effects in this model corresponds to the “biological” (or genotypic) parameterization (*Vitezica et al.* 2013). The dominance coding in the matrix Γ is

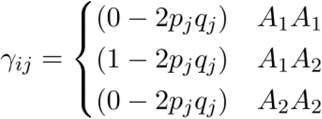

As a result the effects *a* and *d* do not correspond to allele substitution and dominance deviation effects directly, but the sum of variance components still equals the 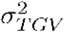 and allele substitution effects can be recovered as *α* = *a* + *d*(*q* − *p*) in order to predict *g_BV_* (Vitezica *et al.* 2013; Varona *et al.* 2018; Werner *et al.* 2020).

### Empirical assessment of the accuracy predicting means, variances, covariances and usefulnesses in cassava crosses

Since 2012, the Next Generation Cassava Breeding project (http://www.nextgencassava.org) has implemented GS in African and Latin American breeding programs (de Oliveira *et al.* 2012; Ly *et al.* 2013; Wolfe *et al.* 2017). Cassava breeding programs are well-poised to adopt *G_MS_* if, in addition to prediction of means, variances and covariances can be accurately predicted.

#### Cassava data: pedigree, genetic map and phased haplotypes

We chose a publicly-available, previously-published pedigree, genetic map and phased marker-dataset as the best starting point for our analysis (Chan *et al.* 2019); https://www.biorxiv.org/content/10.1101/794339v1.full). The pedigree and germplasm chosen represent parents and offspring from the first three cycles of GS conducted at the International Institute of Tropical Agriculture (IITA). These germplasm and selections have been described previously (Wolfe *et al.* 2016a; b, 2017, 2019; Rabbi *et al.* 2017, 2020).

Chan *et al.* (2019) implemented a number of procedures to ensure the quality of the data. First, technical replications of the original genotyping-by-sequencing (GBS) were validated with BIGRED (Chan *et al.* 2018) and reads were combined to reduce missingness and increase read-depth-per-sample. Next, a multi-pass analysis using the pedigree-validation software, AlphaAssign (Whalen *et al.* 2018) was used to ensure only relationships supported by the data were assumed downstream. Genotypes were called using validated pedigree information and were used as input for phasing and imputation. Pedigree-guided imputation and phasing was accomplished by SHAPEIT2/duoHMM (O’Connell *et al.* 2014). Finally, the authors constructed a genetic linkage map based on crossover events observed in the dataset. We accessed the pedigree, genetic map and haplotypes from: http://ftp.cassavabase.org/manuscripts/Chan_et_al_2019/. We restricted our analysis to only the 3199 individuals comprising 462 full-sibling families (and their parents), in which both parents were validated/known.

#### Cassava data: traits, trials and selection indices

We chose four focal cassava traits: dry matter percentage (DM), fresh root yield in natural-log tons-per-hectare (logFYLD), season-wide mean cassava mosaic disease severity (1-5 scale; MCMDS) and total carotenoids by color chart (1-8 scale; TCHART). These traits include both polygenic (DM and logFYLD) and mono/oligenic architectures (MCMDS, TCHART). Two of the traits are known to have important dominance variance (logFYLD and MCMDS), while DM has been shown to be largely additive (Wolfe *et al.* 2016a; b).

From these traits, we composed two hypothetical selection indices, which represent two real and disparate breeding goals **(Table S01)**. Both indices target increased DM and logFYLD and reduced MCMDS. We refer to the first index as the “Standard SI” (or StdSI) as it emphasizes yield and disease resistance in a white-fleshed background. The second index “Biofortification SI” (or BiofortSI) focuses on breaking a historically negative genetic correlation between DM and carotenoid content by weighting most heavily the combination of yellow-flesh (high TCHART) and high DM. We note that the pedigree analyzed here has been selected for the equivalent of the StdSI. For this reason, our population and analyses should not be considered as representative or definitive regarding biofortification breeding goals. We started with unscaled, non-economic weights and scaled them by dividing by the standard deviation of phenotypic BLUPs (see below) for each trait **(Table S01)**.

We used pre-adjusted phenotypes, namely, de-regressed BLUPs as input for our downstream analyses. The field trial data used span from 2013-2019 and are available directly from www.cassavabase.org. The download, quality control, formatting and mixed-model analysis that produced the BLUPs are fully documented and reproducible here: https://wolfemd.github.io/IITA_2019GS/. The BLUPs produced and used in this study of cross variance prediction were originally used for GS conducted during summer 2019. The entire raw IITA trial download was too large for GitHub and is therefore stored here: http://ftp.cassavabase.org/marnin_datasets/NGC_BigData/.

#### Parent-wise cross-validation scheme

We devised a cross-validation scheme that: (1) allowed measurement of the accuracy of predicting means, variances and covariances in previously unobserved crosses, and (2) enabled us to distinguish accuracy predicting BV from TGV.

First, define a vector, P of the parents listed in the pedigree. Define also a second vector **C** listing the genotypes (clones) in the pedigree, including the parents (*P* ⊂ *C*).

We conducted five replications of the following procedure:

1. Define parent-wise cross-validation folds: randomly assign the parents in **P** into *k*-folds. We chose *k* = 5 folds or about 42 of 209 parents in **P** per fold (defined as 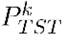, the list of “test” parents in the *k*th-fold.
2. For each of the *k*-folds (set of 42 “test” parents), divide the clones vector **C** into two mutually exclusive sets: “training” (*C_TRN_* and “validation” (*C_VLD_*). From the set *C_TRN_*, we exclude all descendants (offspring, grandchildren, great grandchildren, etc.) of 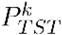. Include the 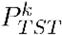 themselves (phenotyping the parents before predicting their offspring) and any remaining non-descendents. Define *C_VLD_* simply as the set difference between **C** and *C_TRN_*.
3. Estimate marker effects independently by fitting mixed-models (see section below for further details) to *C_VLD_* and *C_TRN_* corresponding to each 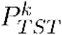.
4. For each 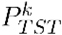, define the set of crosses to predict, 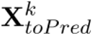 to include any of the 462 actual families (sire-dam pairs) in the pedigree, in which the 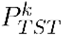 were involved. By construction, the real family members that have been observed for each of the 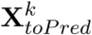 were excluded from the model used to get marker effects for *C_VLD_* and included in the model for Cv w. Predict the means, variances and covariances for each focal trait in each cross, 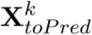 using the *C_TRN_* marker effects only.
5. For each family in 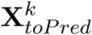, using all existing family members, compute the sample means, variances and covariances for **GEBV** and **GETGV** as predicted by the *C_VLD_* marker effects.
6. Calculate the accuracy of prediction for each mean 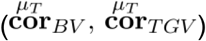, variance 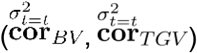 and covariance 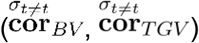 in terms of both **BV** and **TGV**. For 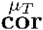 we used the Pearson correlation between predicted and observed mean **GEBV/GETGV**. For 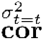 and 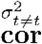 only families with greater than two members were able to be included, and we weighted the correlation between the predicted and sample (co)variance of **GEBV/GETGV** according to the family size (R **package**::*function* **psych**::*cor.wt*). For sake of comparison, we also include accuracies where predicted values are correlated to phenotypic (rather than genomic-predicted) BLUPs, e.g. 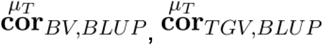, etc.

The cross-validation scheme is numerically summarized in **Tables S02** (see also **Table S03-05)**.

#### Multi-trait Bayesian ridge regressions

We used the multi-trait Bayesian ridge regression (MtBRR) implemented in the development version of the BGLR R package (https://github.com/gdlc/BGLR-R), which is itself a direct port of the model implemented in the package MTM (de los Campos and Grüneberg 2016). The MtBRR models marker effects as being drawn from a multivariate-normal distribution with mean effects of 0 for each trait and variance-covariance parameters jointly estimated from the posterior distribution of the Gibbs chain. We ran each chain for 30000 iterations, discarded the first 5000 as burn-in and thinned to every 5th sample.

We used de-regressed BLUPs as responses in each model to match the approach used for GS (described here, for example: (Wolfe *et al.* 2017)), but BGLR does not currently support weighted observations in the multi-trait model.

We fit the following MtBRR models: (1) additive-only (**A**), (2) additive plus dominance (**ClassicAD**) and (3) additive plus directional dominance (**DirDomAD**).

For each model, we fit an MtBRR to each *C_TRN_* and *C_VLD_* as described above. In addition, we analyzed the entire population (“All” samples) and the component genetic groups, which are: **GG** (or **C0**; the original progenitors chosen from a population known as the “Genetic Gain”), **TMS13**, **TMS14** and **TMS15**, which represent the offspring from 2013 (C1), 2014 (C2) and 2015 (C3), respectively.

#### Predicting cross means, variances and usefulnesses

We predicted cross means using the posterior mean marker effects.

We made variance predictions with the posterior mean marker effects, which we refer to as the variance of posterior means (or **VPM**). Lehermeier *et al.* (2017a; b) used instead the posterior mean variance (**PMV**), which is effectively the mean of the variances predicted by each MCMC-sample of marker effects (see eqns. 7-10 in that study).

The **PMV** is expected to be less biased compared to the **VPM** but is considerably more computationally intensive. Moreover, **PMV** requires the on-disk storage of massive posterior marker-effects arrays. Nevertheless, we computed the **PMV** for each family and compared results to **VPM** to consider whether the faster method is sufficient.

We then computed selection index means and variances using the predicted (and observed) means, variances and covariances of the component traits, and the index weights, given in Table S01.

#### Realized selection intensities (measuring post-cross selection)

We used **GEBV** and **GETGV** based on test-set marker-effects to compute observed (or realized) usefulness criteria i.e. *UC_parent_* and *UC_variety_* and measure prediction accuracy as follows. For *C_VLD_*, we computed the mean **GEBV** of family members who were themselves later used as parents. We computed *UC_variety_* at each **VDP** stage (CET, PYT, AYT, UYT) but focused mostly on the penultimate, advanced yield trial (AYT, e.g. 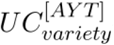). We computed stage-specific *UC_variety_* as the mean **GETGV** of family members advanced to each stage of the **VDP.**

In order to combine predicted means and variances into usefulness criteria, i.e. 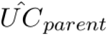 and 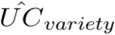, we first calculated the realized intensity of within-family selection (*i_RS_* and *i_VDP_*). For *i_VDP_* we used the proportion of family members who themselves appear in the pedigree as parents. For the *i_VDP_* we used the raw plot-basis data to compute the proportion of clones from each family with at least one plot in each of the aforementioned **VDP** stages as of July 2019. We computed standardized selection intensity in R using *i* = *dnorm*(*qnorm*(1 − *propSel*})/*propSel*, where *propSel* is the proportion selected.

#### Exploratory analysis: predictions of previously untested crosses

We conducted a prediction exercise evaluating the interest of possible future crosses compared to those previously made in terms of additive and total merit, i.e. *UC_parent_* and *UC_variety_*. We predicted the means and variances of all possible pairwise matings between the union of 209 parents already used and the 100 clones with top rank on the StdSI, of which only 3 overlapped(*N*=306 parents). This resulted in 47,083 crosses to predict. We used marker-effects from the full-model (all clones included). We predicted means, variances and covariances for all four traits and subsequently used these to compute StdSI and BiofortSI means and variances.

### Data availability, reproducibility and *predCrossVar* R package

The full repository for this study including all data and output is located at http://ftp.cassavabase.org/manuscripts/Wolfe_et_al_2020. The repository, minus large data files, can be found at https://github.com/wolfemd/PredictOutbredCrossVar/. We used Rmarkdown and the R package **workflowr** (https://github.com/jdblischak/workflowr, version 1.6.2) to document our empirical analysis in a fully reproducible (https://github.com/wolfemd/PredictOutbredCrossVar/) and documented HTML website (https://wolfemd.github.io/PredictOutbredCrossVar/). Finally, we implemented the core functions for multi-trait prediction of outbred cross variances including additive and dominance effects in an R package **predCrossVar** (https://wolfemd.github.io/predCrossVar/, https://github.com/wolfemd/predCrossVar) and used it in the aforementioned analyses.

## Results

Results along with code generating summaries, figures and related tables are also available as part of the **workflowr** R markdown website (Results, Figures, Supplementary Figures, Supplementary Tables).

### Pedigree and Germplasm

There were 3199 individuals in 462 families, derived from 209 parents in our pedigree. Parents were used an average of 31 (median 16, range 1-256) times as male and/or female parents in the pedigree. The mean family size was 7 (median 4, range 1-72). The average percent homozygosity was 0.84 (range 0.76-0.93) across the 3199 pedigree members (computed over 33,370 variable SNP; Table S14). As expected for a population under recurrent selection, the homozygosity rate increased with each generation with C0, C1, C2, and C3 having homozygosity rates of 0.826, 0.835, 0.838, and 0.839, respectively (Figure S01).

### Cross-validation Scheme

Across the 5 replications of 5-fold cross-validation, the average number of clones was 1833 (range 1245-2323) for training sets and 1494 (range 1003-2081) for testing sets. The 25 training-testing pairs set up an average of 167 (range 143-204) crosses to predict (**Table S02**).

### BLUPs and Selection Indices

The correlation between the two SI (StdSI and BiofortSI; **Table S01**) based on *i.i.d.* (non-genomic) BLUPs of component traits was 0.43 (Figure S02). The correlation between DM and TCHART BLUPs was −0.29.

### Accuracy of family mean prediction

On average, across traits, the accuracy of predicting family-mean TGV 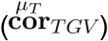 was lower than for the mean BV 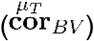 by −0.04 for the ClassicAD model, but essentially the same for the DirDom model. For yield (logFYLD), family 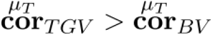 by 0.01 in the ClassicAD model and by 0.13 for the DirDom model **(Figure 1, Table S10)**. The DirDom and ClassicAD models had similar accuracy for 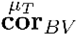 but DirDom gave higher 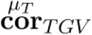 by 0.05 on average and up to 0.11 higher for logFYLD. Note that accuracy is higher for BiofortSI compared to StdSI. This makes sense given that BiofortSI emphasizes DM and TCHART, which have higher accuracy than logFYLD and MCMDS.

**Figure 1.**
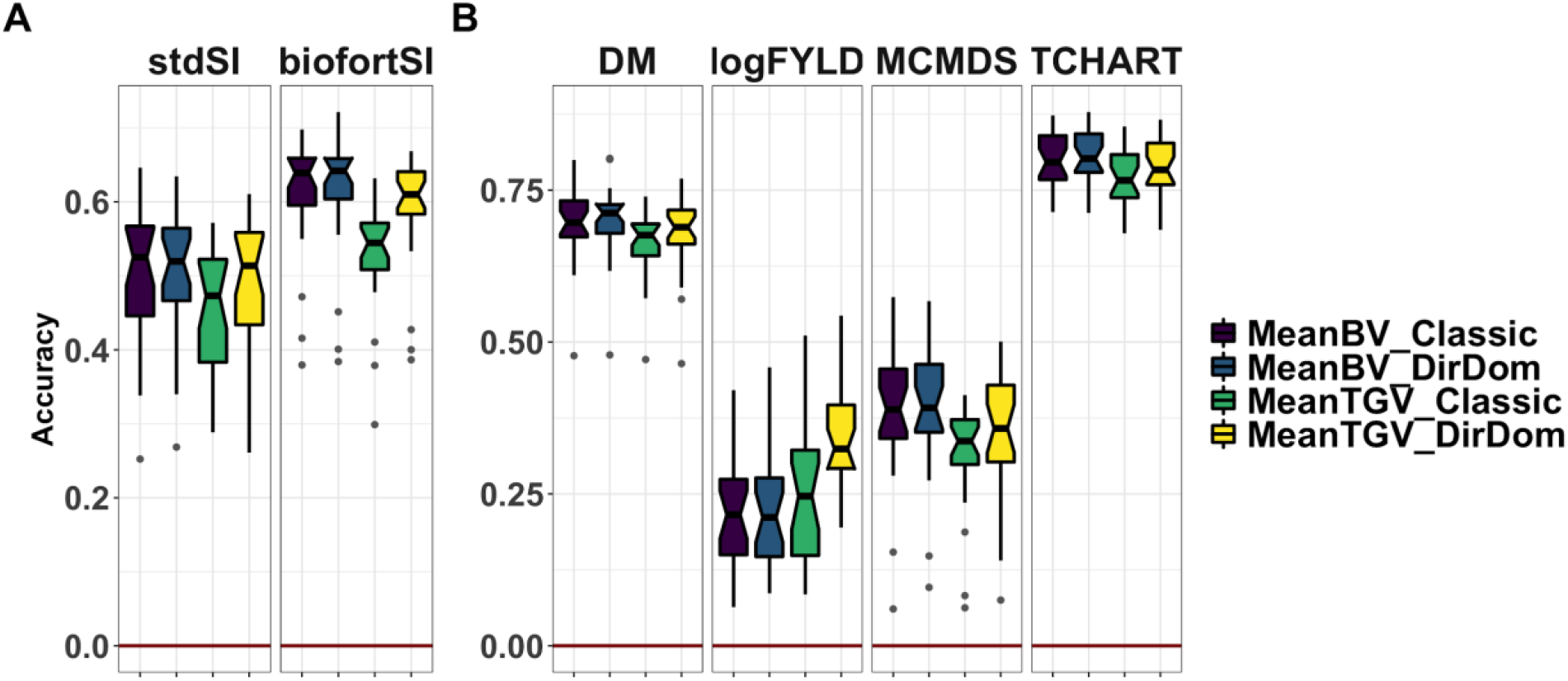
Accuracy predicting the family mean of previously untested crosses. Fivefold parent-wise cross-validation estimates of the accuracy predicting the cross means on selection indices (A) and for component traits (B), is summarized in boxplots. Accuracy (y-axis) was measured as the correlation between the predicted and the observed mean **GEBV** or **GETGV**. For each trait, accuracies for four predictions: two prediction types (family mean **BV** vs. **TGV**) times two prediction models (**Classic** vs. **DirDom**).

### Accuracy of within-family variance and covariance prediction

#### PMV vs. VPM

Variances and covariances were predicted with the computationally intensive PMV method. We began by comparing the PMV and VPM results. Ideally, they will be the same in accuracy and provide similar rankings. VPM is much faster and we would prefer to use it, e.g. for the predictions of untested crosses. Across all predicted variance-covariance components and models, the correlation between predictions using the PMV and VPM was very high (mean correlation of 0.98; **Table S07)**.

However, there is a difference in scale between predictions by VPM and PMV (**Figure S06; Table S07)**. PMV gave consistently higher variance predictions and larger absolute covariance |*σ*|, either more negative i.e. DM-TCHART or more positive i.e. MCMDS-TCHART). In terms of prediction accuracy, PMV-based estimates were nearly uniformly lower (mean decrease in acc. −0.07) **(Figure S07; Table S11)**. We proceeded with PMV results, except for the exploratory analyses of predictions of untested crosses (see below), where we saved time/computation and used the VPM.

#### Comparison of accuracy among models, traits and variance components

We consider the accuracy of predicting variances (and subsequently also usefulness criteria) using the “PMV” variance method, GBLUPs as validation-data and family-size-weighted correlations. Most variance prediction accuracy estimates were positive, with a mean weighted correlation of 0.14 (**Figure 2a, Table S11**). Mean accuracy for covariance prediction was lower at 0.09 (**Figure 2b, Table S11**).

**Figure 2.**
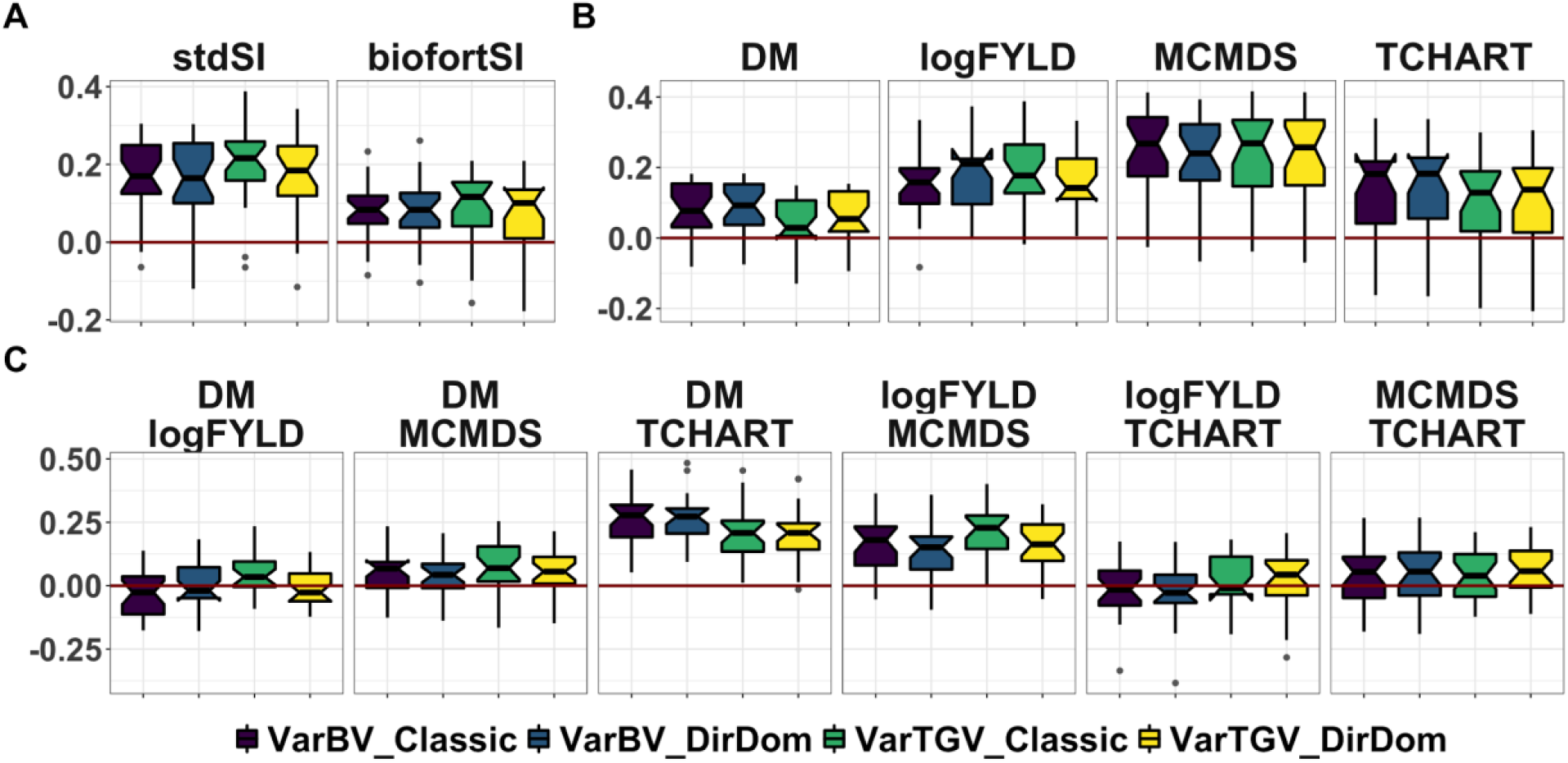
Accuracy predicting the genetic (co)variances of previously untested crosses. Fivefold parent-wise cross-validation estimates of the accuracy predicting the genetic variance of crosses on selection indices (A) and for component trait variances (B) and covariances (C). Accuracy (y-axis) was measured as the correlation between the predicted and the observed (co)variance of **GEBV** or **GETGV**. For each trait (panel), accuracies for four predictions: two prediction types (**VarBV** vs. **VarTGV**) times two prediction models (**Classic** vs. **DirDom**).

In contrast to results for predicting family means, the most accurately predicted trait-variances were MCMDS (mean acc. 0.24) and logFYLD (mean acc. 0.17) while Var(DM), for example, had among the lowest accuracies at 0.07. Interestingly, the DM-TCHART covariance was the most well predicted component (mean acc. 0.24). Accuracy for the selection index variances were intermediate (mean StdSI = 0.18, mean BiofortSI = 0.08) compared to the component traits. Like the accuracy for means on the SI’s, accuracy for variances was related to the accuracy of the component traits. In contrast to predicting cross means on SI’s, for variances, the StdSI > BiofortSI. This makes sense as the StdSI emphasized logFYLD and MCMDS, whose variance were better predicted than those of DM, TCHART and related covariances.

There were, overall, only small differences in accuracy between prediction models (ClassicAD and DirDomAD) and var. components 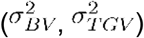. The accuracy of predicting family-(co)variance in TGV averaged higher than in BV for the ClassicAD model, but was the same for DirDom.

On average, the accuracy for StdSI variance was more than twice that of the BiofortSI (0.18 vs. 0.08). For both models and both indices, 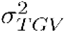 was better predicted than 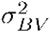.

### Accuracy predicting the usefulness of crosses

The usefulness criteria i.e. *UC_parent_* and *UC_variety_* are predicted by:

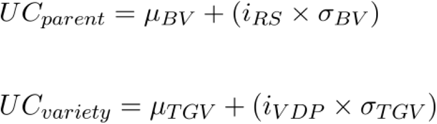

The observed (or realized) **UC** are the mean **GEBV** of family members who were themselves later used as parents. In order to combine predicted means and variances into a **UC**, we first calculated the realized intensity of within-family selection (*i_RS_* and *i_VDP_*) **(Figure S12; Table S13)**. For *UC_parent_* we computed the *i_RS_* based on the proportion of progeny from each family, that themselves later appeared in the pedigree as parents. For *UC_variety_* we computed *i_VDP_* based on the proportion of family-members that had at least one plot at each **VDP** stage (CET, PYT, AYT, UYT). There were 48 families with a mean intensity of 1.59 (mean 2% selected) that themselves had members who were parents in the pedigree. As expected, the number of available families and the proportion selected decreased (increasing selection intensity) from CET to UYT. We chose to focus on the AYT stage, which has 104 families, mean intensity 1.46 (mean 5% selected).

On a per-repeat-fold basis, sample sizes (number of families) with observed usefulness (for measuring prediction accuracy) were limited. For *UC_parent_* there were an average of 17 families (min 9, max 24). For *UC_variety_* the sizes depended on the VDP stage, for the focal stage 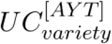, mean number of families was 37 (min 25, max 50). Computing a single accuracy across all repeats, folds *and* models, the *UC_parent_* was more accurately predicted (0.46 stdSI, 0.61 biofortSI) than the 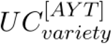 (0.24 stdSI, 0.38 biofortSI). In contrast to predictions of cross variances, the mean UC for the BiofortSI was higher (0.55) compared to the StdSI (0.42), perhaps indicating that the cross-mean dominates the prediction of UC. For the BiofortSI, the ClassicAD and DirDom models had nearly identical accuracy. The DirDom model was slightly more accurate for 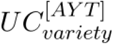, but slightly less so for 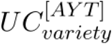 **(Figure 3, Figure S13, TableS12)**. Prediction accuracy for UC was similar when setting a constant intensity of 2.67 **(Figure S13)**.

**Figure 3.**
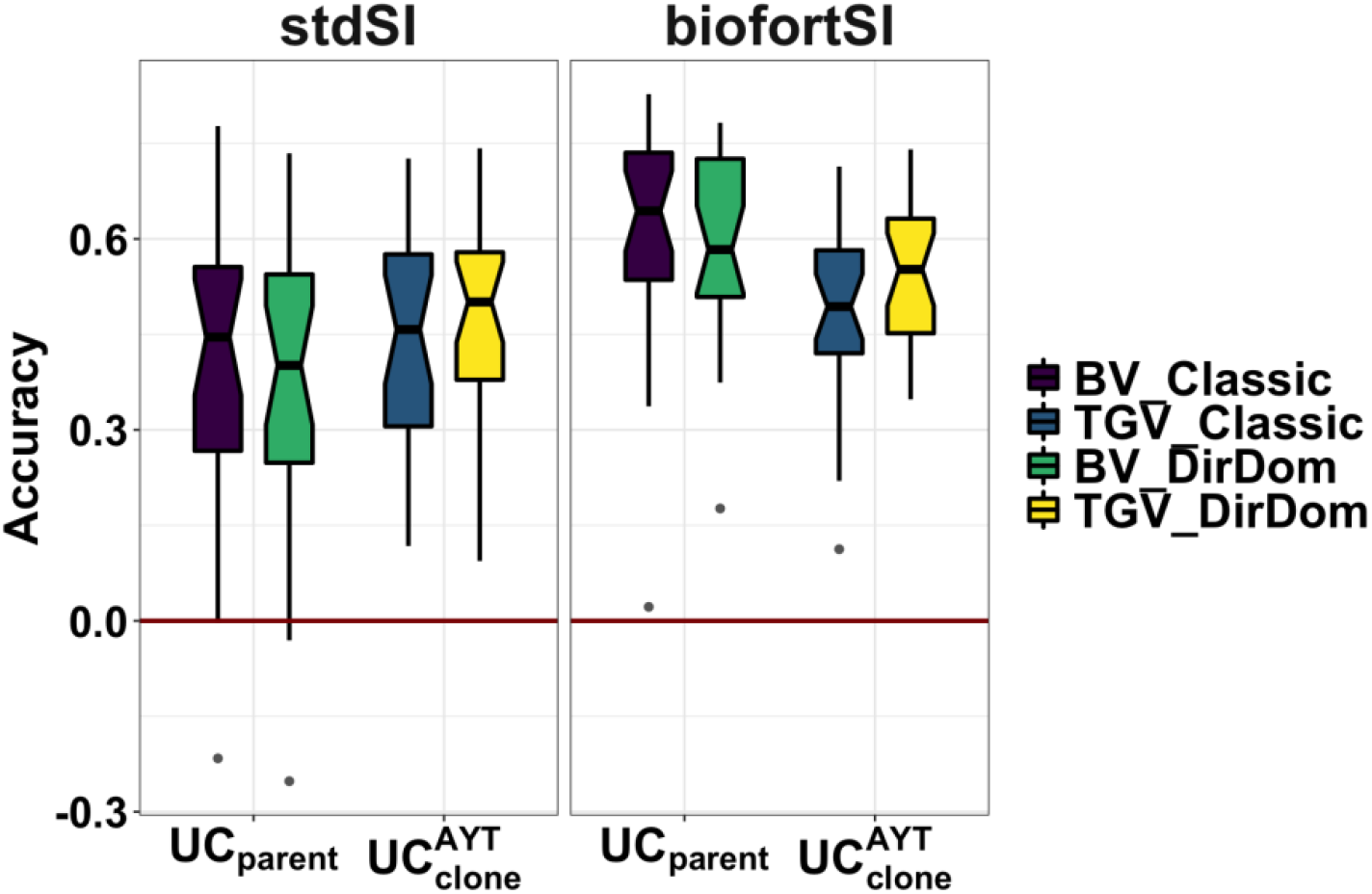
Accuracy predicting the usefulness (the expected mean of future *selected* offspring) of previously untested crosses. Fivefold parent-wise cross-validation estimates of the accuracy predicting the usefulness of crosses on the selection indices (x-axes) is summarized in boxplots. Accuracy (y-axis) was measured as the correlation between the predicted and observed usefulness of crosses for breeding parents (UC_*parent*_) or clones (UC_*variety*_). For each UC (panels), accuracies for four predictions: two selection indices (StdSI and BiofortSI) times two prediction models (**Classic** vs. **DirDom**).

### Population estimates of the importance of dominance variance

Our focus is mainly on distinguishing among crosses, and the accuracy of cross-based predictions. Detailed analysis of the additive-dominance genetic variance-covariance structure in cassava (sub)-populations is an important topic, which we mostly leave for future study. However, we make a brief examination of the genetic variance-covariance estimates associated with the overall population and component genetic groups. We report all variance-covariance estimates in **TableS15** and complete BGLR output in the repository associated with this study. Over all genetic groups analyzed, across traits and SIs, dominance accounted for an average of 34% (range 7-68%) of the genetic variance in the ClassicAD model, and 24% (6-53%) for the DirDomAD model. Across models, dominance was most important (mean 52% of genetic variance) for yield (logFYLD) and least important for DM (mean 22%) and TCHART (mean 13%) **(Figure 4)**. For several estimates, there was an opposing sign between additive and dominance components, i.e. positive additive but negative dominance covariance. For DM-logFYLD there was a tendency for positive dominance but negative additive covariance. For DM-MCMDS, in contrast, the tendency was for negative dominance but positive additive covariance. Both the summary statistics above and **Figure 4** report the estimates according to the **PMV** and used the method for computing variances accounting for LD (method 2, or M2 in (*Lehermeier et al.* 2017a) (but see **Table S15** for method 1 and VPM results).

**Figure 4.**
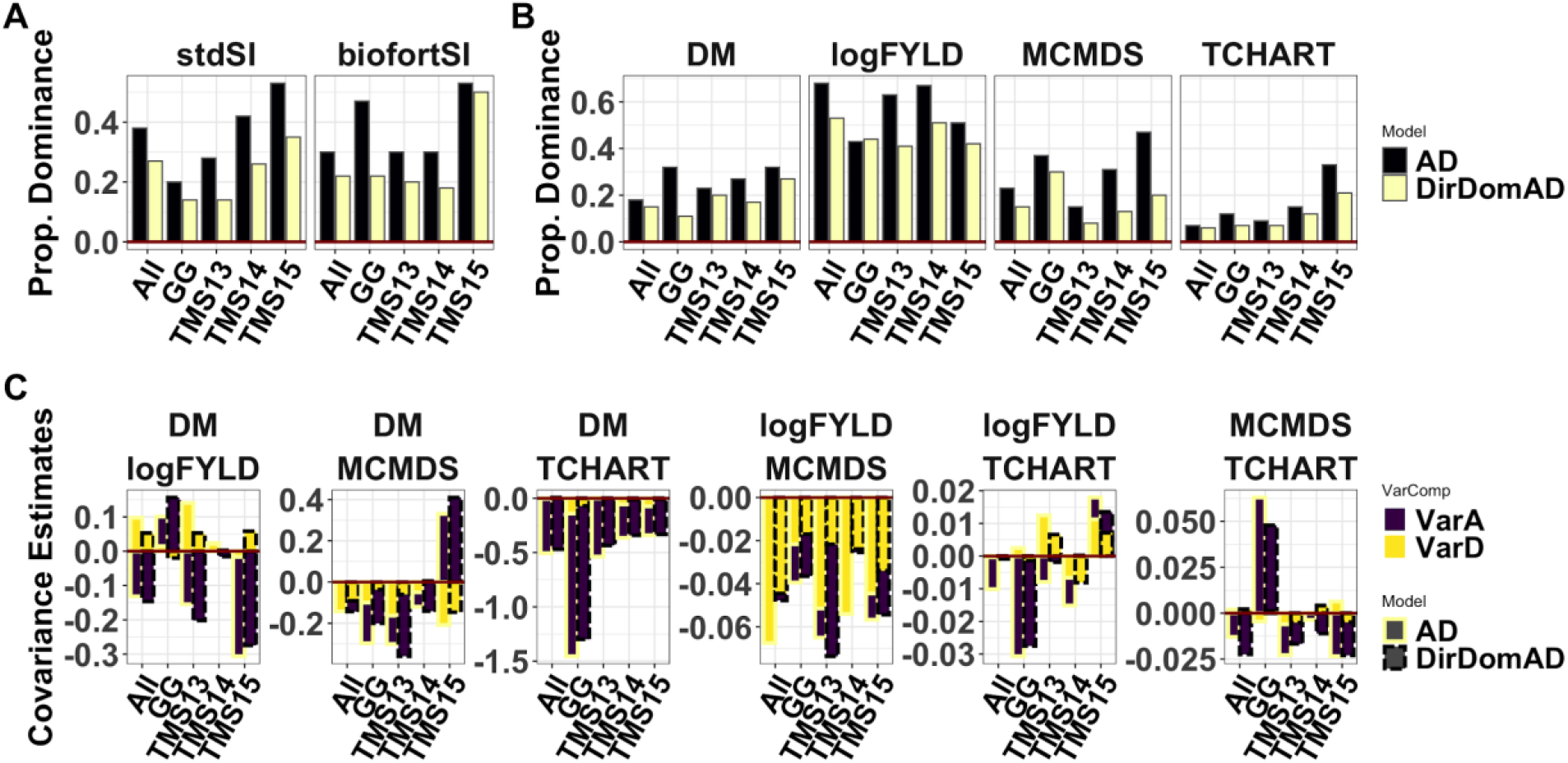
Population-level measures of the importance of dominance genetic effects. The genetic variance estimates from the models fitted to the overall population (“All”) and also to its four genetic groups (x-axis) are presented in these bar plots. Each panel contains results for a trait variance or covariance. For selection indices (A) and component traits (B) the proportion of genetic variance accounted for by dominance is shown on the y-axis. For covariances between component traits (C) the estimates themselves are plotted. For A and B, color distinguishes prediction models (ClassicAD vs. DirDom), whereas for C, color indicates variance component (additive vs. dominance) and models are distinguished by linetype as shown in the legend.

### Population estimates of inbreeding effects

We found mostly consistent and significant (different from zero) effects of inbreeding depression associated especially with logFYLD (*i.e.* mean effect −2.75 log(tons/ha) per genome-wide percent increase in homozygosity across genetic groups, mean effect −3.88 across cross-validation folds), but also DM (−4.82 percent dry matter per unit increase in homozygosity across genetic groups, −7.85 cross-validation) and MCMDS (0.32 worse disease severity per unit increase in homozygosity across genetic groups, 1.27 cross-validation). This corresponds to higher homozygosity being associated with lower DM, lower yield and greater disease severity **(Figure 5, Table S16)**.

**Figure 5.**
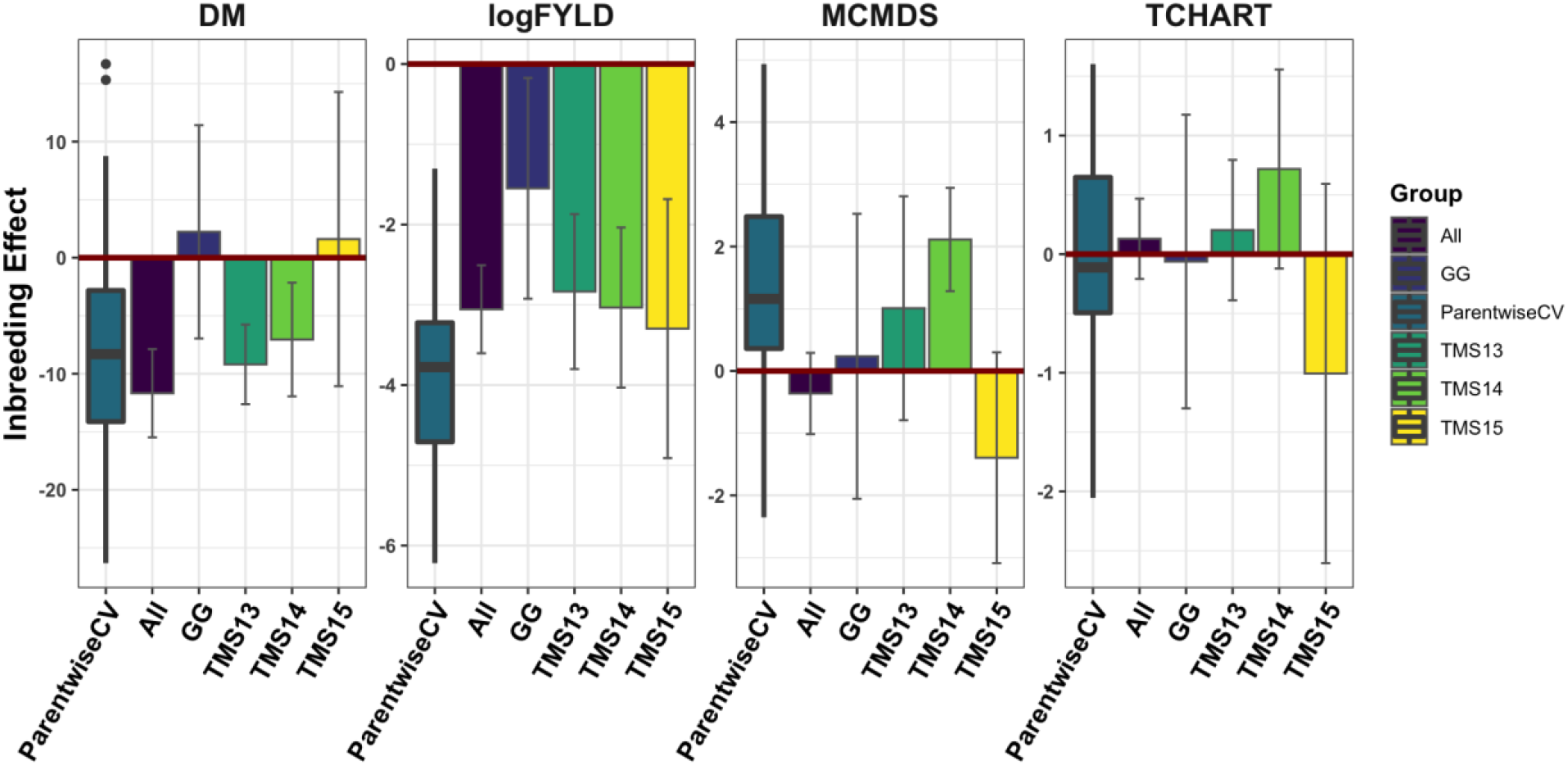
Estimates of the genome-wide effect of inbreeding. For each trait (panels), the fixed-effect for genome-wide proportion of homozygous sites is shown on the y-axis, as estimated by a directional dominance model. For the overall population (“All”) and four genetic groups (“TMS13”, “TMS14”, “TMS15”), the posterior mean estimate and its standard deviation (bars) are shown on the x-axis. For comparison a boxplot showing the distribution of estimates from models fit to parent-wise cross-validation training and validation sets (“ParentwiseCV”) is also shown.

### Exploring predictions about untested crosses

We made 16 predictions (2 SIs x 2 prediction models [ClassicAD, DirDomAD] x 2 selection targets [BV, TGV] x 2 criteria [Mean, UC = Mean + i*SD]) for each of 47,083 possible crosses of 306 parents. We examined the correlation structure among these predictions in order to understand the multivariate decision space they describe **(Figure 6, Figure S14-15).**

**Figure 6.**
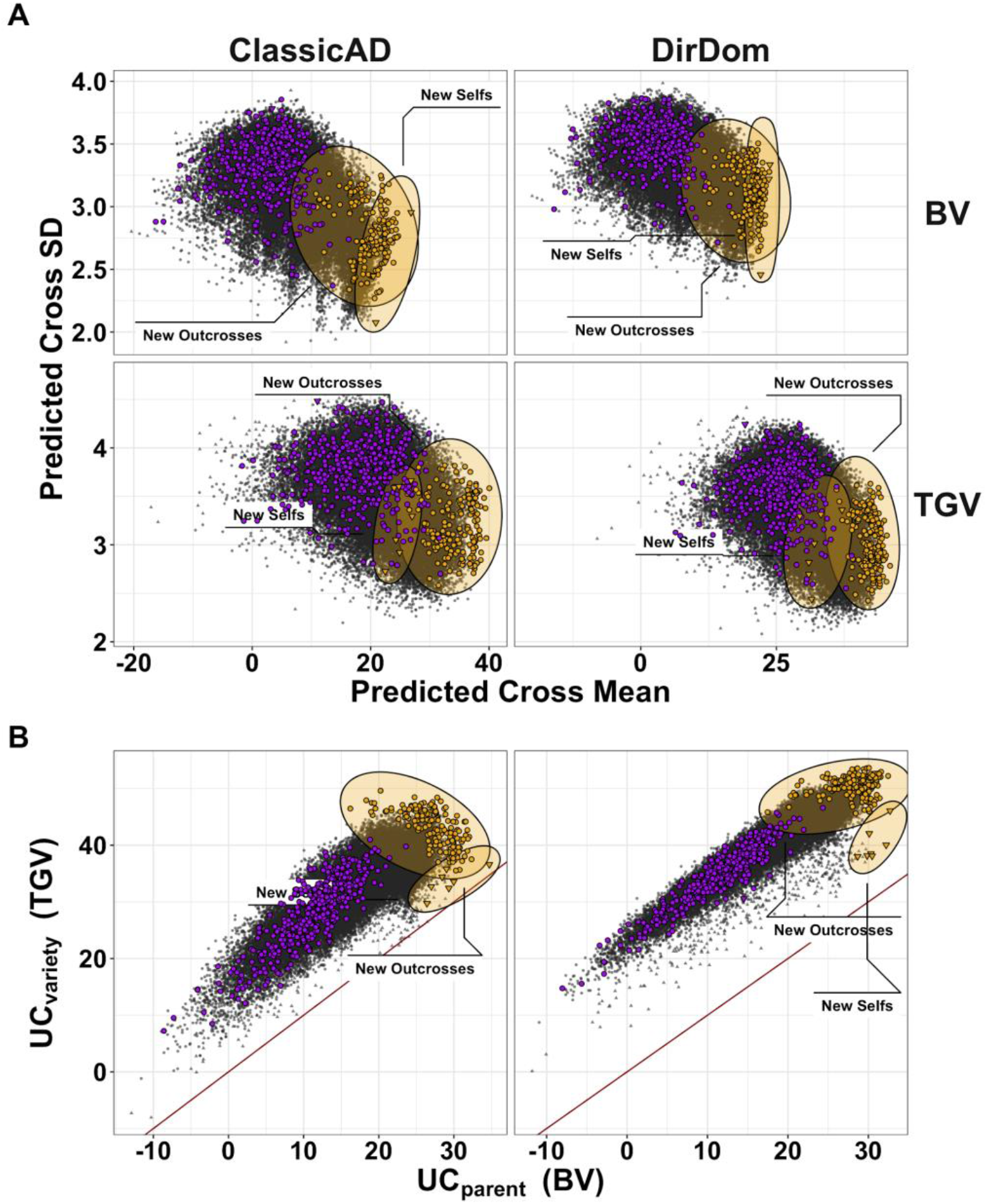
Genomic mate selection criteria for the StdSI predicted for previously untested crosses. We predicted 47,083 crosses among 306 parents. We made eight predictions in total encompassing the 2 prediction models [ClassicAD, DirDomAD] x 2 variance components [BV, TGV] x 2 criteria [Mean, UC = Mean + 2*SD]. Selfs are shown as triangles, outcrosses as circles. For each of the predictions, we took the top 50 ranked crosses and then selected the union of crosses selected by at least one metric for n= 190 “New Crosses”. In each panel, the 190 new crosses are highlighted in yellow and distinguished according to their status as self-vs. outcrosses. The 462 crosses previously made are shown in purple to highlight the opportunity for improvement. The predicted cross genetic mean is plotted against the predicted family genetic standard deviation (Sd, *σ*) for breeding value [BV] and total genetic value [TGV] (panel rows) **(A)**. The *UC_parent_* is also plotted against the *UC_variety_* with a red one-to-one line in **B**. Results are shown for the ClassicAD model (left column) and the DirDomAD model (right column) of **A** and **B**.

The two selection indices are (by design) disparate breeding goals. The mean correlation (across models and var. components) between SIs 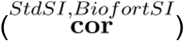 was low for predictions of the family mean (0.18) and lower for the UC (0.12), but high for the SD (0.91). The correlation between predictions made by the ClassicAD and the DirDomAD model 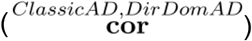 was 0.97 across all predictions. The predictions of BV and TGV were strongly correlated (average 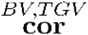 for predicted cross means = 0.91, for predicted variances = 0.94 and predicted UC = 0.90). The biggest difference (lower correlation) was for the ClassicAD model and the StdSI 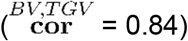.

The predicted cross means and variances had a low, but negative correlation **(Figure 6A)**. Across traits, models and variance components, the average correlation between predicted mean and standard deviation 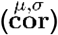 was −0.36. At the standardized intensity of 2.67 (1% selected) the predicted UC was dominated by the mean 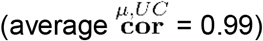) and there was a small negative correlation between variance and UC 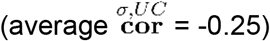.

We wanted to know how selections of crosses-to-make would be affected by our choice of criteria. Separately, for each of the 16 predictions of 47,083 crosses, we selected the top 50 ranked crosses **(TableS19)**. In total, only 310 unique crosses were selected based on at least one of the 16 predictions. Of those, 190 were selected for the StdSI (120 Biofort) and included only 7 (7) self-crosses. No crosses were selected for both SI. None of the selected crosses have previously been tested in the IITA breeding program. We plotted the predicted *μ* vs. the UC **(Figure 6A)** the *UC_parent_* vs. the *UC_variety_* **(Figure 6B)** for both ClassicAD and DirDomAD results. We highlight the 190 unique new crosses proposed and contrast them to the 462 previously made, in order to illustrate that our genomic mate selection criteria propose substantially better crosses. For simplicity, we plotted predictions for StdSI only.

There were 87 parents represented among the 190 “best” crosses for StdSI with a median usage in 3 families each (range 1-169, most popular parent = **TMS13F1095P0013**). Only 42 parents were indicated for the BiofortSI with a median contribution to 5 (range 1-149, most popular parent = **IITA-TMS-IBA011371**) crosses. **Figure 7** breaks down the selections on the StdSI according to model, prediction and variance components as a network where selected parents are nodes and matings are edges. For the StdSI only 52 crosses were selected by both ClassicAD and DirDomAD, 78 and 60 were uniquely selected by each model, respectively. For the BiofortSI 71 crosses were selected by both models and 30 and 19 were unique to ClassicAD and DirDomAD models, respectively. Only 23 of 190 (StdSI) and 37 of 120 crosses (BiofortSI) were selected for *both* BV and TGV.

**Figure 7.**
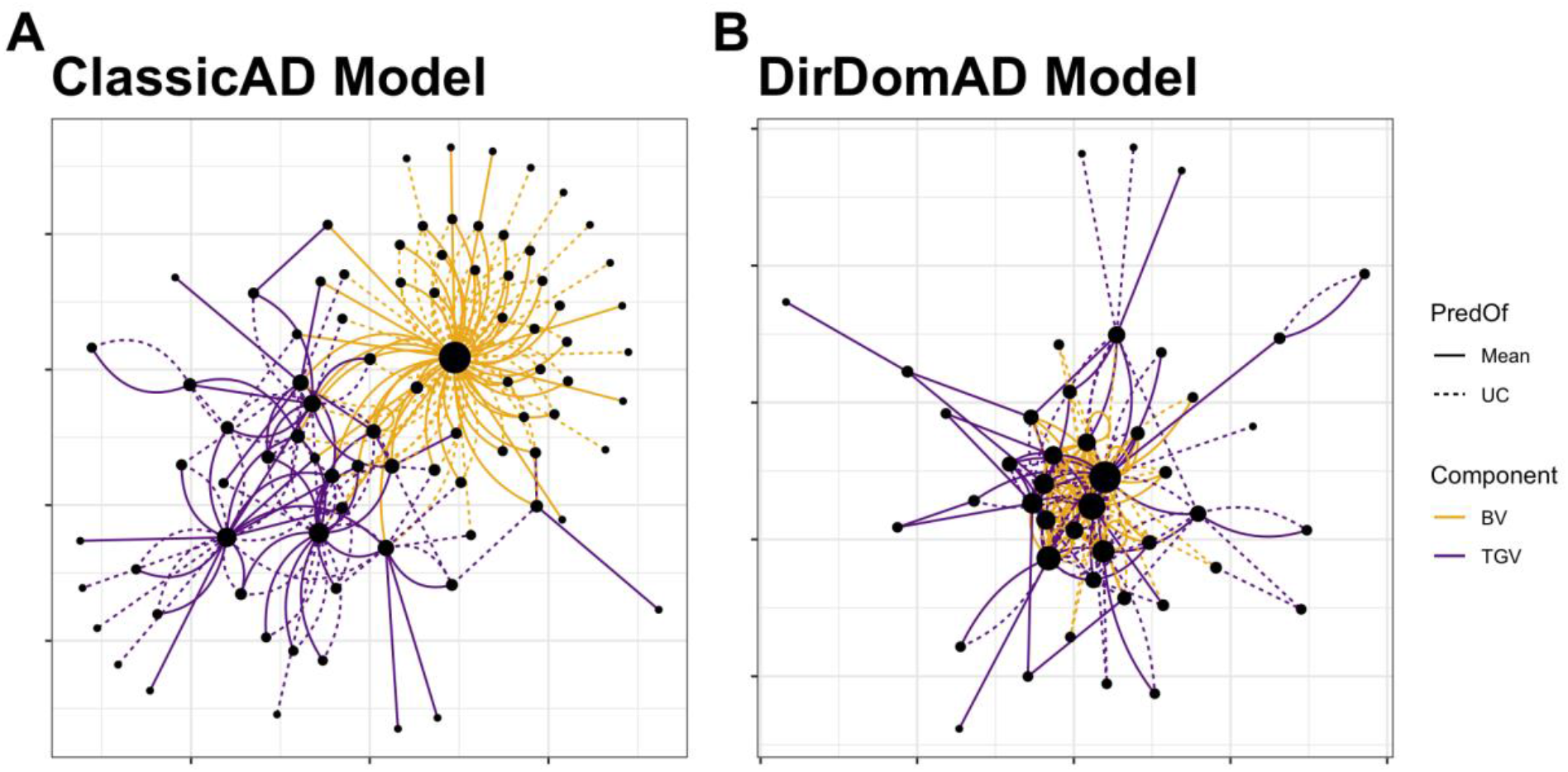
Network plot of selected parents and matings for the StdSI. There were 87 parents and 190 crosses chosen because they were in the top 50 for at least one of eight criteria (2 prediction models [ClassicAD, DirDomAD] x 2 variance components [BV, TGV] x 2 criteria [Mean, UC = Mean + 2*SD]). Parents are shown as nodes, with size proportional to their usage (number of connections). Matings are shown as edges, with linetype distinguishing selection based on Mean (solid) and UC (dashed) and color depicts selection for breeding value, BV (orange) vs. total genetic value, TGV (purple). Selections arising from the ClassicAD model **(A)** and the DirDomAD model **(B)** are shown in panels.

In fact, 28 of 87 parents selected on the StdSI were chosen only for the TGV of crosses and 26 only for BV (Figure 7). For the BiofortSI, no parents were chosen only for BV, but 23 of 42 were only interesting for their TGV. Only 39 crosses for the StdSI (18 for BiofortSI) were selected *only* based on the UC (i.e. selected for their variance *but not* their mean).

So which are the “best” crosses to make? If you use the number of times chosen as a criterion and you don’t break ties, there are 112 unique crosses in the top 50 for the StdSI and 50 for the BiofortSI.

## Discussion

We developed and tested genomic *mate* selection criteria suitable for multi-trait index selection in clonal organisms of arbitrary homozygosity level where the F1 (full-sibling progeny) are of direct interest as future parents and/or cultivars (varieties). We focused on the prediction of the selection index-associated genetic variance of a cross based on the haplotypes of proposed parents, estimates of marker effects and a genetic map (estimates of recombination frequencies between marker loci). We combined the predicted mean and variance of a cross into usefulness criteria (UC) for parent (*UC_parent_*) and variety (*UC_variety_*) development, by predicting the variance in terms of both breeding (*σ_BV_*) and total genetic values *σ_TGV_* = *σ_BV_* + *σ_DD_*.

### Sufficiency of mean and variance prediction accuracy estimates and their implications

Among diploid, outbred, clonally propagated crops, cassava has well developed genomic resources (Rabbi *et al.* 2020) and is entering advanced stages of GS at breeding programs in Africa (Wolfe *et al.* 2017) and Latin America (*de Oliveira *et al.* 2012; Andrade et al.* 2019). Here, we worked with 462 real cassava families of heterogeneous size. We made practical use of the available data in implementing the parent-wise cross-validation scheme. We found that accuracy predicting genetic variances were greater than zero both for component traits *and* selection indices. As theory predicts and all previous studies show, the accuracy of variance (second-moment) prediction was less than that of means (first-moment) (Zhong and Jannink 2007; Osthushenrich *et al.* 2018; Neyhart and Smith 2019).

Prediction accuracies for cassava trait means were similar to previously published estimates (Wolfe *et al.* 2016b, 2017). However, unlike previous studies, our cross-validation scheme allows us to use genomic estimates of BV and TGV as validation data. Rather than correlating predicted means and variances to phenotypes (or *i.i.d.* estimates) of individual lines, we computed the sample means and variances of BV and TGV for each validation family. As a result, and in contrast to previous results, prediction of TGV was less accurate overall compared to BV predictions.

Many factors contribute to achieving optimal accuracy and those factors are well understood in the literature. We focused here on getting an assessment of the overall ability to distinguish crosses with high vs. low genetic variances. Previous studies of variance-prediction accuracy evaluated relatively few families, but with larger size (Yao *et al.* 2018; Osthushenrich *et al.* 2018; Neyhart and Smith 2019). Interestingly, we found that traits with the most accurately predicted variances had less accurately predicted means, including the SIs (mean: StdSI < BiofortSI; variance: StdSI > BiofortSI; **Figure 1, Figure 2**). This does not seem initially explainable by our priors regarding trait genetic architectures; DM and FYLD are both generally considered as polygenic / infinitesimal traits, while MCMDS and TCHART are expected to be closer to mono- or oligogenic (Wolfe *et al.* 2016a; Rabbi *et al.* 2020). Differences in accuracy between mean and variance predictions should instead have to do with the nature of linkage disequilibrium, especially as it affects marker-causal relationships (de Los Campos *et al.* 2015; Lehermeier *et al.* 2017a). Similar to the simulations and empirical results of (Neyhart *et al.* 2019) we found that the DM-TCHART covariance was particularly well predicted, corresponding to hypothesized tight linkage between QTL on chromosome 1 (Rabbi *et al.* 2017); a region known to contain large low-recombination regions of historical introgression from the wild relative *M. glaziovii* (Wolfe *et al.* 2019).

We suggest that the actual accuracy achievable is higher than our estimates, as our validation data itself is subject to the sampling error of our particular pedigree. That error decreases the correlation, biasing all our estimates downward. That is, while our prediction is of a progeny pool of infinite size, our validation sample is finite and heterogeneous in size. An alternative measure of variance prediction accuracy could be achieved through *in silico* simulation of hundreds of offspring per cross, with variance in BV computed using test-set marker effects. However, this strategy would only increase accuracy estimates by decreasing sampling error associated with the validation sample size.

### The importance of non-additive effects and the effect on inbreeding

Non-additive effects are important in cassava, accounting for an average of 34% of genetic variance in this study. Our results are consistent with previous studies that highlight the importance of non-additive effects for fresh root yields but not for dry matter or total carotenoid content (*Wolfe *et al.* 2016b, 2017; Esuma *et al.* 2016; Nduwumuremyi *et al.* 2018; Andrade et al. 2019).* To our knowledge, we are the first to report partitions of additive and dominance genetic *co*variance, though we do not comment on it in detail in this article. We also report the first estimates of genome-wide marker-based directional dominance in cassava. Using the model of (Xiang *et al.* 2016) we found significant and consistent evidence of inbreeding *depression,* for every trait except TCHART, but especially yield. Our results match several previous estimates of inbreeding depression based on field observation of selfed (S1) progeny (Pujol and McKey 2006; Rojas *et al.* 2009; Kawuki *et al.* 2011; de Freitas *et al.* 2016). Theory and data (reviewed in (Kristensen and Sørensen 2005)) indicate that traits more closely associated with fitness (in cassava, this would be traits related to root and stem production, for example) should be more impacted by directional dominance (inbreeding depression). These results also make sense in light of the evidence of deleterious genetic load in cassava (*Ramu et al. 2017)* and balancing selection for heterozygosity in introgression regions (Wolfe *et al.* 2019).

It was also interesting to observe that different crosses were selected (esp. by ClassicAD) for BV and TGV. In fact, the network plot in **Figure 7** shows a pattern of crosses within vs. among particular parents selected for exploiting either BV or TGV. This pattern is suggestive of heterotic groups and warrants future investigation.

### Caveats, limitations and future directions for GMS in outbred, clonal crops

For practicality, it is preferable to predict cross-variances using the variance of posterior means (VPM) rather than the posterior mean variance (PMV). Our results show that the correlation between VPM and PMV predictions is very high *but* the magnitude is different, as is the accuracy estimate. PMV is expected to be less biased and as such, the correct weight (selection intensity) for computing the UC can be chosen relative to the intended proportion of selection (Lehermeier *et al.* 2017a; b; Neyhart and Smith 2019). If any bias is consistent, then re-ranking between PMV and VPM (or REML) predictions of cross variance is not a concern. Either way it is possible that using PMV would increase the apparent importance of the predicted genetic variance when computing the UC.

Other critical considerations for practical implementation include the necessary phasing quality and method. We leveraged a dataset imputed and phased using a validated pedigree (Chan *et al.* 2019); many plant breeding programs may not have suitable pedigree or depth of relationships to enable this. We do not rule out using “standard” population-based imputation and phasing e.g. Browning and Browning (2016). Promising also will be the development of a practical haplotype graph suitable for outbred diploids like cassava (e.g. Zou *et al.* 2020; Jensen *et al.* 2020). In addition, the necessary marker density for accurate prediction should be considered as it has a very significant effect on computational speed.

Several extensions and future directions are of interest moving forward from the current study. We have only addressed dominance, but extensions of variance prediction to include epistasis or even non-linear kernel types should be straightforward (Alves *et al.* 2019). The directional dominance model and its assumption of uncorrelated additive and dominance effects and linear genome-wide effects on phenotype of increasing homozygosity need evaluation (Xiang *et al.* 2018). We note that *many* outbred, clonal crops are actually polyploids. For organisms with such genomes, further developments in recombination mapping, phasing and prediction models will be required, but are expected to be possible. In our study, we focused on trait-associated variance prediction. Considerable development of mate selection criteria has concerned the avoidance of genetic diversity loss *generally,* these are approaches that constrain inbreeding (Kinghorn 2011; Woolliams *et al.* 2015) and are distinct from trait-associated predictions presented here. We note that Allier *et al.* (2019a) recently described prediction of the variance in parental contribution in a family (i.e. the variance in inbreeding level) as a correlated trait, using an extension of the approach for prediction trait-associated variance.

### Conclusions

By providing predictions of the selection-index-associated means *and* variances in arbitrary crosses for additive *and* dominance variances, we provided a suite of genomic *mate* selection criteria suitable for the complexities of a modern (cassava) breeding program. We presented a simple approach for genomic *truncation* mate selection that identifies a profile of crosses collectively interesting because of the predicted merit of their progeny in terms of *μ_BV_, μ_TGV_, σ_BV_* and *μ_TGV_*. Ultimately, crossing plans can be numerically optimized (Akdemir and Sánchez 2016; Gorjanc and Hickey 2018; Akdemir *et al.* 2019; Allier *et al.* 2019b) to consider trait-associated means and variances as well as inbreeding levels, to provide a high degree of control for the management of breeding populations.

## Acknowledgements

We are grateful to the entire Next Generation Cassava Breeding team (www.nextgencassava.org) and especially the International Institute of Tropical Agriculture Cassava Breeding team, so many of whom have contributed to this study in the field, in the lab and beyond. We appreciate Andrew Werner for pointing us towards directional dominance models, and the Jean-Luc Jannink and Mark Sorrells research groups for fruitful discussions and comments along the way. Thanks to Lukas Mueller and Prasad Peteti for data hosting and curation respectively.

## Supplementary Tables

Most Supplementary Tables are included as worksheets in the file **SupplementaryTables.xlsx**. Very large ones are included as separate CSV files.

Table S1: Selection indices. For each trait, the standard deviation of BLUPs, which were divided by “unscaled” index weights for the StdSI and BiofortSI indices to get StdSI and BiofortSI weights used throughout the study.

Table S2: Summary of cross-validation scheme. For each fold of each Rep, the number of parents in the test-set (Ntestparents) is given along with the number of clones in the corresponding training (Ntraintset) and testing (Ntestset) datasets and the number of crosses to predict (NcrossesToPredict).

Table S3: Test-parents. For each fold of each cross-validation repeat, the set of parents whose crosses are to be predicted is listed.

Table S4: Training-Testing partitions of germplasm. For each fold of each repeat, the genotype ID (germplasmName) of all clones in the “trainset” and “testset” are given.

Table S5: Crosses to predict each fold. For each fold of each repeat, the sireID and damID are given for each cross-to-be-predicted.

Table S6: Predicted and observed cross means. For each fold of each repeat, each cross distinguished by a unique pair of sireID and damID is given. The genetic model used (Models A, AD, DirDomAD, DirDomBV), whether the prediction is of mean breeding value (predOf=MeanBV) or mean total genetic value (predOf=MeanTGV), the trait (BiofortSI or StdSI), type of observation (ValidationData: GBLUPs or iidBLUPs) and corresponding prediction (predMean) and observations (obsMean) are shown.

Table S7: Predicted cross variances. All predictions of cross-variance from the cross-validation scheme are detailed. For each fold of each repeat and each unique cross (sireID-damID). Both variances (Trait1 ==Trait2) and co-variances (Trait1!=Trait2) are given. The genetic model used (Models A, AD, DirDomAD, DirDomBV), the variance component being predict (VarA or VarD), along with the number of segregating SNPs in the family (Nsegsnps) and the time taken in seconds for computation, per family (totcomputetime) are given. The predictions based on the variance of posterior means (VPM) and the posterior mean variances (PMV) are both shown.

Table S8: Predicted and observed cross variances. For each fold of each repeat, each cross distinguished by a unique pair of sireID and damID is given. The genetic model used (Models A, AD, DirDomAD, DirDomBV), whether the prediction is of family variance in breeding value (predOf=VarBV) or variance in total genetic value (predOf=VarTGV), the trait (BiofortSI or StdSI), type of observation (ValidationData: GBLUPs or iidBLUPs) and corresponding prediction (predVar) and observations (obsVar) are shown. The predictions are based on either only the variance of posterior means (VarMethod=VPM) or the posterior mean variances (VarMethod=PMV). The family size (number of genotyped offspring, FamSize) or number of offspring with direct phenotypes (Nobs) are used (CorrWeight) to weight the correlation between observed and predicted family variances.

Table S9: Predicted and observed UC. For each fold of each repeat, each cross distinguished by a unique pair of sireID and damID is given. The predicted usefulness criterion (predUC) was computed as the predMean + realIntensity\*predSD, where predMean is the predicted family mean and predSD is the predicted genetic standard deviation. The genetic model used (Models A, AD, DirDomAD, DirDomBV), whether the prediction is of family variance in breeding value (predOf=VarBV) or variance in total genetic value (predOf=VarTGV), the trait (BiofortSI or StdSI) and corresponding prediction (predUC) and observations (obsUC) are shown. The family size (number of genotyped offspring, FamSize) is shown along with the realized selection intensity (realIntensity) for each selection stage in the breeding pipeline (Parent, CET, PYT, AYT, UYT) and also a constant intensity value (Stage=ConstIntensity).

Table S10: Accuracy of predicting the mean. For each fold of each repeat, the accuracy predicting family means (Accuracy) is given. The genetic model used (Models A, AD, DirDomAD, DirDomBV), whether the prediction is of mean breeding value (predOf=MeanBV) or mean total genetic value (predOf=MeanTGV), the trait (BiofortSI or StdSI), type of observation (ValidationData: GBLUPs or iidBLUPs) are shown.

Table S11: Accuracy of predicting the variances. For each fold of each repeat the estimated accuracy of predicting family variances is given. Accuracy was computed the correlation between predicted and observed variance, either weighted by family size (AccuracyWtCor) or not (AccuracyCor). The genetic model used (Models A, AD, DirDomAD, DirDomBV), whether the prediction is of family variance in breeding value (predOf=VarBV) or variance in total genetic value (predOf=VarTGV), the trait (BiofortSI or StdSI), type of observation (ValidationData: GBLUPs or iidBLUPs) are shown. The predictions are based on either only the variance of posterior means (VarMethod=VPM) or the posterior mean variances (VarMethod=PMV).

Table S12: Accuracy predicting the usefulness criteria. For each fold of each repeat the estimated accuracy of predicting family usefulness criteria is given. Accuracy was computed as the correlation between predicted UC and observed UC (mean of selected offspring), either weighted by family size (AccuracyWtCor) or not (AccuracyCor). The genetic model used (Models A, AD, DirDomAD, DirDomBV), whether the prediction is of UC in breeding value (predOf=VarBV) or UC in total genetic value (predOf=VarTGV), the trait (BiofortSI or StdSI), type of observation (ValidationData: GBLUPs or iidBLUPs) are shown.

Table S13: Realized within-cross selection metrics. Table summarizing measurements made of selection within each cross (unique sireID-damID). Summaries included: family size (FamSize), number and proportion of members used as parents, mean GEBV and GETGV of top 1% of each family, for each selection index (BiofortSI, StdSI), proportion of each family that has been phenotyped and past each stage of the breeding pipeline (past CET, PYT, AYT) and finally the corresponding realized intensity of selection for each stage.

Table S14: Proportion homozygous per clone. Genome-wide proportion of SNPs that are homozygous, for each clone in the study.

Table S15: Variance estimates for genetic groups. Summary of the population-level genetic variance estimates in each genetic group (GG, TMS13, TMS14, TMS15), for each genetic model (A, AD, DirDomA, DirDomAD), each variance (Trait1==Trait2) and covariance (Trait1 !=Trait2). The estimates are computed both based on the variance of posterior means (VarMethod=VPM) and the posterior mean variances (VarMethod=PMV). The “Method” refers to whether linkage disequilibrium is accounted for (M2) or not (M1).

Table S16: Directional dominance effects estimates. For each trait in each genetic group and each fold of each cross-validation repeat of the directional dominance model, the posterior mean and standard deviation of the effect of genome-wide homozygosity is given.

Table S17: Predictions of untested crosses. Compiled predictions of 47,083 possible crosses of 306 parents. Predictions were made either with the classic additive-plus-dominance (ClassicAD) or the directional dominance (DirDomAD) model. Whether the cross is a self and/or has previously been made is indicated along with the number of segregating SNPs expected in the family. The predicted mean, standard deviation and usefulness in terms of breeding values (BV) and total genetic values is given.

Table S18: Long-form table of predictions about untested crosses. Compiled predictions of 47,083 possible crosses of 306 parents. Predictions were made either with the classic additive-plus-dominance (ClassicAD) or the directional dominance (DirDomAD) model. Whether the cross is a self and/or has previously been made is indicated along with the number of segregating SNPs expected in the family. The predicted mean, standard deviation and usefulness in terms of breeding values (BV) and total genetic values is given.

Table S19: Top 50 crosses selected by each criterion. For each of the 16 predictions of 47,083 crosses, select the top 50 ranked crosses.

## Supplementary Figures

Figure S01: Boxplot of the genome-wide proportion of homozygous SNPs in each of four genetic groups comprising the study pedigree.

Figure S02: Correlations among BLUPs (including Selection Indices). (A) StdSI vs. BiofortSI computed from i.i.d. BLUPs. (B) Heatmap of the correlation among BLUPs for each of four component traits and two derived selection indices.

Figure S03: Scatterplot comparing accuracies for family means using different validation-data.

Figure S04: Boxplots to show that Accuracies per trait-fold-rep-Model do not re-rank much whether using the iid or the GBLUPs as validation data.

Figure S05: Accuracy predicting family means - GBLUP vs. iid-BLUPs as validation data. Fivefold parent-wise cross-validation estimates of the accuracy predicting the cross means on selection indices and for component traits (x-axis), summarized in boxplots. Accuracy (y-axis) was measured as the correlation between the predicted and the observed mean GEBV or GETGV. For each trait, accuracies for four predictions: two prediction types (family mean BV vs. TGV) times two prediction models (Classic vs. DirDom). Validation data (GBLUPs vs. iidBLUPs) are shown in two horizontal panels.

Figure S06: The difference between PMV and VPM for variance and covariance predictions. Each boxplot shows the posterior mean variance (PMV) minus the variance of posterior means (VPM) based prediction of cross variances and covariances. Each panel is a variance or covariance. Each boxplot shows either an additive or dominance variance from one of the genetic models (x-axis).

Figure S07: The difference between PMV and VPM in terms of prediction accuracy. Each boxplot shows the posterior mean variance (PMV) minus the variance of posterior means (VPM) based estimate of prediction accuracy for cross variances and covariances. Each panel is a variance or covariance. Each boxplot shows either an additive or dominance variance from one of the genetic models (x-axis).

Figure S08: Scatterplot comparing accuracies for family variance and covariance prediction using different validation-data.

Figure S09: Plot of variance-prediction accuracy to show whether re-ranking occurrs according to the choice of validation data (x-axis), GBLUPs vs. i.i.d. BLUPs.

Figure S10: Accuracy predicting family variances - GBLUP vs. iid-BLUPs as validation data. Fivefold parent-wise cross-validation estimates of the accuracy predicting (A) genetic variances and (B) covariances. Selection indices and component trait variances are shown on the x-axis. Accuracy (y-axis) was measured as the weighted correlation between the predicted and the observed (co)variance of GEBV or GETGV. For each trait (panel), accuracies for four predictions: two prediction types (VarBV vs. VarTGV) times two prediction models (Classic vs. DirDom). Validation data (GBLUPs vs. iidBLUPs) are shown in horizontal panels.

Figure S11: Accuracy predicting family (co)variances - Weighted vs. Unweighted Correlation. Fivefold parent-wise cross-validation estimates of the accuracy predicting (A) genetic variances and (B) covariances. Selection indices and component trait variances are shown on the x-axis. Accuracy (y-axis) was measured as the (weighted or unweighted) correlation between the predicted and the observed (co)variance of GEBV or GETGV. For each trait (panel), accuracies for four predictions: two prediction types (VarBV vs. VarTGV) times two prediction models (Classic vs. DirDom). Weighted vs. Unweighted Correlations as accuracy estimates are shown in horizontal panels.

Figure S12: Realized selection intensities: measuring post-cross selection. Boxplots showing (A) the proportion of each family selected and (B) the standardized selection intensity for each stage of the breeding pipeline, in each genetic group.

Figure S13: Accuracy Predicting Usefulness Criteria - All comparisons. Accuracy predicting the usefulness (the expected mean of future selected offspring) of previously untested crosses. Fivefold parent-wise cross-validation estimates of the accuracy predicting the usefulness of crosses on the selection indices (x-axes) is summarized in boxplots. Accuracy (y-axis) was measured as the correlation between the predicted and observed usefulness of crosses for each breeding pipeline stage as well as at a constant selection intensity (x-axis). For each UC (panels), accuracies for four predictions: two selection indices (StdSI and BiofortSI) times two prediction models (Classic vs. DirDom).

Figure S14: Correlation matrix for predictions on the StdSI. Heatmap of the correlations between predictions of mean, standard deviation, and usefulness in terms of BV and TGV, for both the classic and directional dominance model. Predictions were made for 47,083 possible pairwise crosses of 306 parents.

Figure S15: Correlation matrix for predictions on the BiofortSI. Heatmap of the correlations between predictions of mean, standard deviation, and usefulness in terms of BV and TGV, for both the classic and directional dominance model. Predictions were made for 47,083 possible pairwise crosses of 306 parents.

## Author Contributions

Designed the research: MW, JLJ.

Performed the research: MW.

Contributed analyses: AWC.

Contributed data: PK, IYR.

Wrote the paper: MW.

Contributed to drafting the manuscript: AWC, PK, IYR, JLJ

All authors gave final approval for the publication.

## References

Akdemir D., and J. I. Sánchez, 2016 Efficient Breeding by Genomic Mating. Front. Genet. 7:210.

Akdemir D., W. Beavis, R. Fritsche-Neto, A. K. Singh, and J. Isidro-Sánchez, 2019 Multi-objective optimized genomic breeding strategies for sustainable food improvement. Heredity 122: 672–683.

Allier A., L. Moreau, A. Charcosset, S. Teyssèdre, and C. Lehermeier, 2019a Usefulness Criterion and Post-selection Parental Contributions in Multi-parental Crosses: Application to Polygenic Trait Introgression. G3 9: 1469–1479.

Allier A., C. Lehermeier, A. Charcosset, L. Moreau, and S. Teyssèdre, 2019b Improving Short-and Long-Term Genetic Gain by Accounting for Within-Family Variance in Optimal Cross-Selection. Front. Genet. 10: 1006.

Alves F. C., Í. S. C. Granato, G. Galli, D. H. Lyra, R. Fritsche-Neto, et al., 2019 Bayesian analysis and prediction of hybrid performance. Plant Methods 15: 14.

Andrade L. R. B. de, M. B. E. Sousa, E. J. Oliveira, M. D. V. de Resende, and C. F. Azevedo, 2019 Cassava yield traits predicted by genomic selection methods. PLoS One 14:e0224920.

Bijma P., Y. C. J. Wientjes, and M. P. L. Calus, 2020 Breeding Top Genotypes and Accelerating Response to Recurrent Selection by Selecting Parents with Greater Gametic Variance. Genetics 214: 91–107.

Browning B. L., and S. R. Browning, 2016 Genotype Imputation with Millions of Reference Samples. Am. J. Hum. Genet. 98: 116–126.

Campos G. de los, and A. Grüneberg, 2016 Mtm (multiple-trait model) package [www document]. URL http://quantgen.github.io/MTM/vignette.html (accessed 10. 25. 17).

Chan A. W., A. L. Williams, and J.-L. Jannink, 2018 A statistical framework for detecting mislabeled and contaminated samples using shallow-depth sequence data. BMC Bioinformatics 19: 478.

Chan A. W., A. L. Williams, and J.-L. Jannink, 2019 Sexual dimorphism and the effect of wild introgressions on recombination in Manihot esculenta. 794339.

Elias A. A., I. Rabbi, P. Kulakow, and J.-L. Jannink, 2018 Improving Genomic Prediction in Cassava Field Experiments Using Spatial Analysis. G3: Genes|Genomes|Genetics 8: 53–62.

Esuma W., R. S. Kawuki, L. Herselman, and M. T. Labuschagne, 2016 Diallel analysis of provitamin A carotenoid and dry matter content in cassava (Manihot esculenta Crantz).Breed. Sci. 66: 627–635.

Falconer D. S., and T. F. C. Mackay, 1996 Introduction to quantitative genetics (4th edn). Pearson, Prentice Hall, Harlow.

Freitas J. P. X. de, V. da Silva Santos, and E. J. de Oliveira, 2016 Inbreeding depression in cassava for productive traits. Euphytica 209: 137–145.

Gaynor R. C., G. Gorjanc, A. R. Bentley, E. S. Ober, P. Howell, et al., 2017 A two-part strategy for using genomic selection to develop inbred lines. Crop Sci. 57: 2372–2386.

Gemenet D. C., and A. Khan, 2017 Opportunities and Challenges to Implementing Genomic Selection in Clonally Propagated Crops, pp. 185–198 in Genomic Selection for Crop Improvement: New Molecular Breeding Strategies for Crop Improvement, edited by Varshney R. K., Roorkiwal M., Sorrells M. E. Springer International Publishing, Cham.

Gorjanc G., R. Chris Gaynor, and J. M. Hickey, 2018 Optimal cross selection for long-term genetic gain in two-part programs with rapid recurrent genomic selection. Theoretical and Applied Genetics 131: 1953–1966.

Gorjanc G., and J. M. Hickey, 2018 AlphaMate: a program for optimizing selection, maintenance of diversity and mate allocation in breeding programs. Bioinformatics 34: 3408–3411.

Heffner E. L., M. E. Sorrells, and J.-L. Jannink, 2009 Genomic Selection for Crop Improvement. Crop Sci. 49: 1–12.

Heslot N., H.-P. Yang, M. E. Sorrells, and J.-L. Jannink, 2012 Genomic Selection in Plant Breeding: A Comparison of Models. Crop Sci. 52: 146–160.

Hickey J. M., Implementing Genomic Selection in CGIAR Breeding Programs Workshop Participants, T. Chiurugwi, I. Mackay, and W. Powell, 2017 Genomic prediction unifies animal and plant breeding programs to form platforms for biological discovery. Nature Genetics 49: 1297–1303.

Jannink J.-L., A. J. Lorenz, and H. Iwata, 2010 Genomic selection in plant breeding: from theory to practice. Brief. Funct. Genomics 9: 166–177.

Jensen S. E., J. R. Charles, K. Muleta, P. J. Bradbury, T. Casstevens, et al., 2020 A sorghum practical haplotype graph facilitates genome-wide imputation and cost-effective genomic prediction. Plant Genome 13: 1687.

Kawuki R., E. Nuwamanya, L. Herselman, M. E. Ferguson, and Others, 2011 Segregation of selected agronomic traits in six S1 cassava families. J. Plant Breed. Crop Sci. 3: 154–160.

Kinghorn B. P., 2011 An algorithm for efficient constrained mate selection. Genet. Sel. Evol. 43:4.

Kristensen T. N., and A. C. Sørensen, 2005 Inbreeding – lessons from animal breeding, evolutionary biology and conservation genetics. Animal Science 80: 121–133.

Lehermeier C., G. de Los Campos, V. Wimmer, and C.-C. Schön, 2017a Genomic variance estimates: With or without disequilibrium covariances? J. Anim. Breed. Genet. 134: 232–241.

Lehermeier C., S. Teyssèdre, and C.-C. Schön, 2017b Genetic Gain Increases by Applying the Usefulness Criterion with Improved Variance Prediction in Selection of Crosses. Genetics 207: 1651–1661.

Los Campos G. de, D. Sorensen, and D. Gianola, 2015 Genomic heritability: what is it? PLoS Genet. 11: e1005048.

Ly D., M. Hamblin, I. Rabbi, G. Melaku, M. Bakare, et al., 2013 Relatedness and genotype×environment interaction affect prediction accuracies in genomic selection: a study in cassava. Crop Sci. 53: 1312–1325.

Lynch M., and B. Walsh, 1998 Genetics and analysis of quantitative traits. Sinauer Sunderland, MA.

Nduwumuremyi A., R. Melis, P. Shanahan, and A. Theodore, 2018 Genetic inheritance of pulp colour and selected traits of cassava (Manihot esculenta Crantz) at early generation selection. J. Sci. Food Agric. 98: 3190–3197.

Neyhart J. L., A. J. Lorenz, and K. P. Smith, 2019 Multi-Trait Improvement by Predicting Genetic Correlations in Breeding Crosses. G3 (Bethesda, Md.) 9: 3153–3165.

Neyhart J. L., and K. P. Smith, 2019 Validating Genomewide Predictions of Genetic Variance in a Contemporary Breeding Program. Crop Sci. 59: 1062–1072.

O’Connell J., D. Gurdasani, O. Delaneau, N. Pirastu, S. Ulivi, et al., 2014 A general approach for haplotype phasing across the full spectrum of relatedness. PLoS Genet. 10: e1004234.

Okeke U. G., D. Akdemir, I. Rabbi, P. Kulakow, and J.-L. Jannink, 2017 Accuracies of univariate and multivariate genomic prediction models in African cassava. Genet. Sel. Evol. 49: 88.

Oliveira E. J. de, M. D. V. de Resende, V. da Silva Santos, C. F. Ferreira, G. A. F. Oliveira, et al., 2012 Genome-wide selection in cassava. Euphytica 187: 263–276.

Osthushenrich T., M. Frisch, C. Zenke-Philippi, H. Jaiser, M. Spiller, et al., 2018 Prediction of Means and Variances of Crosses With Genome-Wide Marker Effects in Barley. Front. Plant Sci. 9: 1899.

Ozimati A., R. Kawuki, W. Esuma, I. S. Kayondo, M. Wolfe, et al., 2018 Training Population Optimization for Prediction of Cassava Brown Streak Disease Resistance in West African Clones. G3 8: 3903–3913.

Pujol B., and D. McKey, 2006 Size asymmetry in intraspecific competition and the density-dependence of inbreeding depression in a natural plant population: a case study in cassava (Manihot esculenta Crantz, Euphorbiaceae). J. Evol. Biol. 19: 85–96.

Rabbi I. Y., L. I. Udoh, M. Wolfe, E. Y. Parkes, M. A. Gedil, et al., 2017 Genome-Wide Association Mapping of Correlated Traits in Cassava: Dry Matter and Total Carotenoid Content. Plant Genome 10: 1–14.

Rabbi I. Y., S. I. Kayondo, G. Bauchet, M. Yusuf, C. I. Aghogho, et al., 2020 Genome-wide association analysis reveals new insights into the genetic architecture of defensive, agro-morphological and quality-related traits in cassava. Plant Mol. Biol. https://doi.org/10.1007/s11103-020-01038-3

Ramu P., W. Esuma, R. Kawuki, I. Y. Rabbi, C. Egesi, et al., 2017 Cassava haplotype map highlights fixation of deleterious mutations during clonal propagation. Nat. Genet. 49: 959–963.

Rojas M. C., J. C. Pérez, H. Ceballos, D. Baena, N. Morante, et al., 2009 Analysis of inbreeding depression in eight S1 cassava families. Crop Sci. 49: 543–548.

Santantonio N., and K. Robbins, 2020 A hybrid optimal contribution approach to drive short-term gains while maintaining long-term sustainability in a modern plant breeding program. 2020.01.08.899039.

Santos D. J. A., J. B. Cole, T. J. Lawlor Jr, P. M. VanRaden, H. Tonhati, et al., 2019 Variance of gametic diversity and its application in selection programs. J. Dairy Sci. 102: 5279–5294.

Schnell F. W., and H. F. Utz, 1976 F1 Leistung und Elternwahl in der Zuchtung von Selbstbefruchtern. Ber Arbeitstag Arbeitsgem Saatzuchtleiter.

Segelke D., F. Reinhardt, Z. Liu, and G. Thaller, 2014 Prediction of expected genetic variation within groups of offspring for innovative mating schemes. Genet. Sel. Evol. 46: 42.

Toro M. A., and L. Varona, 2010 A note on mate allocation for dominance handling in genomic selection. Genet. Sel. Evol. 42: 33.

Varona L., A. Legarra, M. A. Toro, and Z. G. Vitezica, 2018 Non-additive Effects in Genomic Selection. Frontiers in Genetics 9.

Vitezica Z. G., L. Varona, and A. Legarra, 2013 On the additive and dominant variance and covariance of individuals within the genomic selection scope. Genetics 195: 1223–1230.

Werner C. R., R. Chris Gaynor, D. J. Sargent, A. Lillo, G. Gorjanc, et al., 2020 Genomic selection strategies for clonally propagated crops. 2020.06.15.152017.

Whalen A., G. Gorjanc, and J. M. Hickey, 2018 Parentage assignment with genotyping-by-sequencing data. J. Anim. Breed. Genet. 228999.

Wolfe M. D., I. Y. Rabbi, C. Egesi, M. Hamblin, R. Kawuki, et al., 2016a Genome-wide association and prediction reveals genetic architecture of cassava mosaic disease resistance and prospects for rapid genetic improvement. Plant Genome 9: 1–13.

Wolfe M., P. Kulakow, I. Y. Rabbi, and J.-L. Jannink, 2016b Marker-based estimates reveal significant non-additive effects in clonally propagated cassava (Manihot esculenta):implications for the prediction of total genetic value and the selection of varieties. G3:Genes, Genomes, Genetics 6: 3497–3506.

Wolfe M. D., D. P. Del Carpio, O. Alabi, L. C. Ezenwaka, U. N. Ikeogu, et al., 2017 Prospects for Genomic Selection in Cassava Breeding. Plant Genome 10. https://doi.org/10.3835/plantgenome2017.03.0015

Wolfe M. D., G. J. Bauchet, A. W. Chan, R. Lozano, P. Ramu, et al., 2019 Historical Introgressions from a Wild Relative of Modern Cassava Improved Important Traits and May Be Under Balancing Selection. Genetics 213: 1237–1253.

Woolliams J. A., P. Berg, B. S. Dagnachew, and T. H. E. Meuwissen, 2015 Genetic contributions and their optimization. J. Anim. Breed. Genet. 132: 89–99.

Xiang T., O. F. Christensen, Z. G. Vitezica, and A. Legarra, 2016 Genomic evaluation by including dominance effects and inbreeding depression for purebred and crossbred performance with an application in pigs. Genet. Sel. Evol. 48: 92.

Xiang T., O. F. Christensen, Z. G. Vitezica, and A. Legarra, 2018 Genomic Model with Correlation Between Additive and Dominance Effects. Genetics 209: 711–723.

Yao J., D. Zhao, X. Chen, Y. Zhang, and J. Wang, 2018 Use of genomic selection and breeding simulation in cross prediction for improvement of yield and quality in wheat (Triticum aestivum L.). The Crop Journal 6: 353–365.

Yonis B. O., D. P. del Carpio, M. Wolfe, J.-L. Jannink, P. Kulakow, et al., Improving root characterisation for genomic prediction in cassava

Zhong S., and J.-L. Jannink, 2007 Using quantitative trait loci results to discriminate among crosses on the basis of their progeny mean and variance. Genetics 177: 567–576.

Zou C., A. Karn, B. Reisch, A. Nguyen, Y. Sun, et al., 2020 Haplotyping the Vitis collinear core genome with rhAmpSeq improves marker transferability in a diverse genus. Nat. Commun. 11: 413.

